# DNA polymerase Lambda is anchored within the NHEJ synaptic complex via Ku70/80

**DOI:** 10.1101/2024.08.12.607588

**Authors:** Philippe Frit, Himani Amin, Sayma Zahid, Nadia Barboule, Chloe Hall, Gurdip Matharu, Steven W. Hardwick, Jeanne Chauvat, Sébastien Britton, Dima Y. Chirgadze, Virginie Ropars, Jean-Baptiste Charbonnier, Patrick Calsou, Amanda K. Chaplin

## Abstract

Non-homologous end joining (NHEJ) is the predominant pathway by which double-strand DNA breaks (DSBs) are repaired in mammals. To enable final break closure, various NHEJ end-processing factors respond to the chemistry of the damaged DNA ends. Amongst these factors is DNA polymerase lambda (Pol λ), a member of the Pol X family. How members of the Pol X family engage with the NHEJ complex is unknown. Here, we present cryo-EM structures of Pol λ in complex with the Ku70/80 DSB sensor whilst engaged with the DNA-PK holoenzyme in a long-range synaptic complex. These structures reveal a specific interaction site between Ku70/80 and the Pol λ BRCT domain. The functionality of this interaction is assessed by generating point mutations on either side of the Pol λ BRCT:Ku70/80 interface. Using these mutants in two orthogonal assays in cells (live protein recruitment at biphoton laser-damaged nuclear sites and transfection with an original gap-filling reporter plasmid) defines the molecular basis and essentiality of the BRCT domain for the recruitment and activity of the Pol λ within the NHEJ complex. Ultimately, these data explain the role of this interaction in cell survival to DSBs. Finally, we propose a unified model for the interaction of the three Pol X family members bearing BRCT domains with the same site of Ku70/80.

## Introduction

Non-homologous end joining (NHEJ) is the predominant pathway in which double strand DNA (dsDNA) breaks are repaired in mammals. Central to the process of NHEJ are large, multi-protein complexes formed by the canonical proteins DNA-dependent protein kinase catalytic subunit (DNA-PKcs), the heterodimer of Ku70/80, DNA ligase IV (LigIV), X-ray repair cross-complementing protein 4 (XRCC4) and XRCC4-like factor (XLF) ^1^. In addition, PAXX (Paralog of XRCC4 and XLF) is an accessory NHEJ protein identified more recently which has functional redundancy with XLF in the NHEJ mechanism ^2,3^. Although this repair pathway is a complex process, it is generally considered to proceed via three major steps; DNA end recognition, DNA end processing and finally DNA ligation ^4^.

The initial step of dsDNA break recognition relies predominantly on the Ku70/80 heterodimer, which engages the DNA ends and subsequently recruits the DNA-PKcs kinase, together forming the DNA-PK holoenzyme ^5^. Recent cryo-electron microscopy (cryo-EM) structures of DNA-PK assemblies revealed how DNA substrates can be positioned for efficient synapsis via dimerization of this large enzyme. Intriguingly, DNA-PK has two distinct mechanisms to form a pre-synaptic complex, corresponding to alternate structural dimers of DNA-PK. It was first proposed that DNA-PK can synapse the broken DNA ends using a dimer of DNA-PK mediated by the C-terminal region of Ku80 (herein termed Ku80-mediated dimer) ^6^. Soon after, an alternative DNA-PK dimer complex was described, which can form upon addition of XRCC4, LigIV and XLF ^2,7,8^. In this second complex, the DNA-PK dimer is crucially bridged by XLF (herein termed XLF-mediated dimer). In both assemblies, the distance between the damaged DNA ends is identical at 115 Å. This distance is not close enough for DNA ligation to proceed without further rearrangement of the assembly, therefore both dimeric assemblies have been referred to as long-range synaptic complexes (LRC). We have shown recently that PAXX can replace XLF for bridging the Ku80-mediated DNA-PK dimer through binding to Ku70 but does not replace XLF in the XLF mediated dimer ^2^.

The second step in the NHEJ mechanism is end processing and is crucial if the DNA ends cannot be readily ligated. Different end processing factors can be recruited to either trim or extend the damaged ends via the action of DNA nucleases or polymerases, as detailed below.

The final step of the repair mechanism involves ligation of the phosphodiester backbone with 5’-phosphate and 3’-hydroxyl moieties by LigIV, facilitated by XLF and XRCC4. To enable ligation, the LRC is thought to transition to a short-range complex (SRC) following autophosphorylation and removal of DNA-PKcs ^9^. In this SRC, the damaged DNA ends are in much closer proximity to each other and thereby are primed for DNA ligation. A cryo-EM structure of the SRC has been solved which shows the catalytic domain of LigIV interacting directly with the DNA^7^. While structural studies have significantly improved understanding of the NHEJ mechanism by illuminating the architecture of several complexes central to the process, how the end processing proteins, in particular polymerases, interact with these complexes has remained an enigma.

The end processing factors known to be implicated in NHEJ include nucleases such as Artemis which can trim DNA nucleotides, and polymerases which can be used for end filling or extension. Recent structural data has revealed how Artemis can engage with a DNA-PK monomer via its N-terminal nuclease domain and is positioned between the N-HEAT (Huntingtin, Elongation Factor 3, A subunit of protein phosphatase 2A, Target of Rapamycin/TOR) and M-HEAT domain of DNA-PKcs ^10^. To date however, the structural mechanism of interaction between DNA polymerases and the NHEJ machinery has not been defined. The polymerases involved in NHEJ belong to the DNA Polymerases X family and catalyse the addition of nucleotides at the 3’-OH end of DNA. In mammals, three polymerases, namely Pol λ, Pol μ and terminal deoxynucleotidyl transferase (TdT), account for the majority of DNA synthesis during NHEJ, with the latter mainly involved in V(D)J recombination ^11,12 13^. These polymerases share a common domain organisation which consists of a breast cancer gene 1 (BRCA1) C-terminal (BRCT) region at the N-terminus followed by a C-terminal catalytic domain **(Figure S1A)**. The catalytic region facilitates the protein:DNA interaction, thus enabling the function of the enzyme. The chemical nature of the DNA ends is a key determinant of the polymerase activity, with each of the three enzymes having varying degrees of template-dependency. Pol λ appears to display strong activity on DNA ends that have a paired primer terminus ^14^. Furthermore, due to its accuracy, it has been proposed that Pol λ is given priority for gap filling in most cell types ^15^. The BRCT domains are thought to direct the polymerases to sites of DNA damage via interactions with complexes comprising DNA and Ku70/80, or larger assemblies including the NHEJ proteins DNA-PKcs, LigIV and XRCC4 ^16,17^. While the structure of the BRCT and catalytic domain of these polymerases have been solved in isolation ^16,18^, how these domains engage with each other, and larger NHEJ complexes has remained unclear.

In order to elucidate how the polymerases interact with NHEJ machineries, we collected cryo-EM data of the Ku80 mediated dimer of DNA-PK in complex with Pol λ. From our structural data we identify the specific binding region of the BRCT domain of Pol λ with the bridge region of Ku70/80 within DNA-PK complex. Using mutagenesis studies in cells on both Pol λ and Ku70/80, we identified key residues that are critical for Pol λ anchoring in NHEJ complexes, efficient gap-filling activity and ultimately cell survival to DSBs. Finally, we propose that all members of the Polymerase X family share a conserved recognition site on the Ku70/80 heterodimer.

## Results

### Cryo-EM structure of Pol λ bound to the Ku80-mediated DNA-PK dimer

In order to understand how Pol λ interacts with the NHEJ Long Range Complex (LRC), cryo-EM data was collected on a sample containing DNA-PKcs, Ku70/80, DNA, XRCC4, LigIV, PAXX and Pol λ to allow formation of the Ku80-mediated dimer of DNA-PK. Following extensive particle classification, a consensus map was obtained of the Ku80-mediated dimer of DNA-PK. The particles comprising this initial map were further characterized to generate maps of the Ku80-mediated dimer with and without XRCC4 and DNA LigIV (thereafter termed LX4) engaged **(Figure 1, S2-4)**. At this stage, it was apparent that there were no large areas of additional density that could accommodate the full-length Pol λ protein, when compared to previous maps of this DNA-PK dimer. However, a small region of additional density was apparent at the bridge that defines the thinner part of the Ku70/80 ring region **(Figure S5)**. To focus specifically on this region, particles were re-extracted in smaller boxes corresponding to the monomeric size of DNA-PK, and maps were generated of single protomers from within the dimeric assemblies (herein termed half-dimers) with and without LX4 bound.

**Figure 1:**
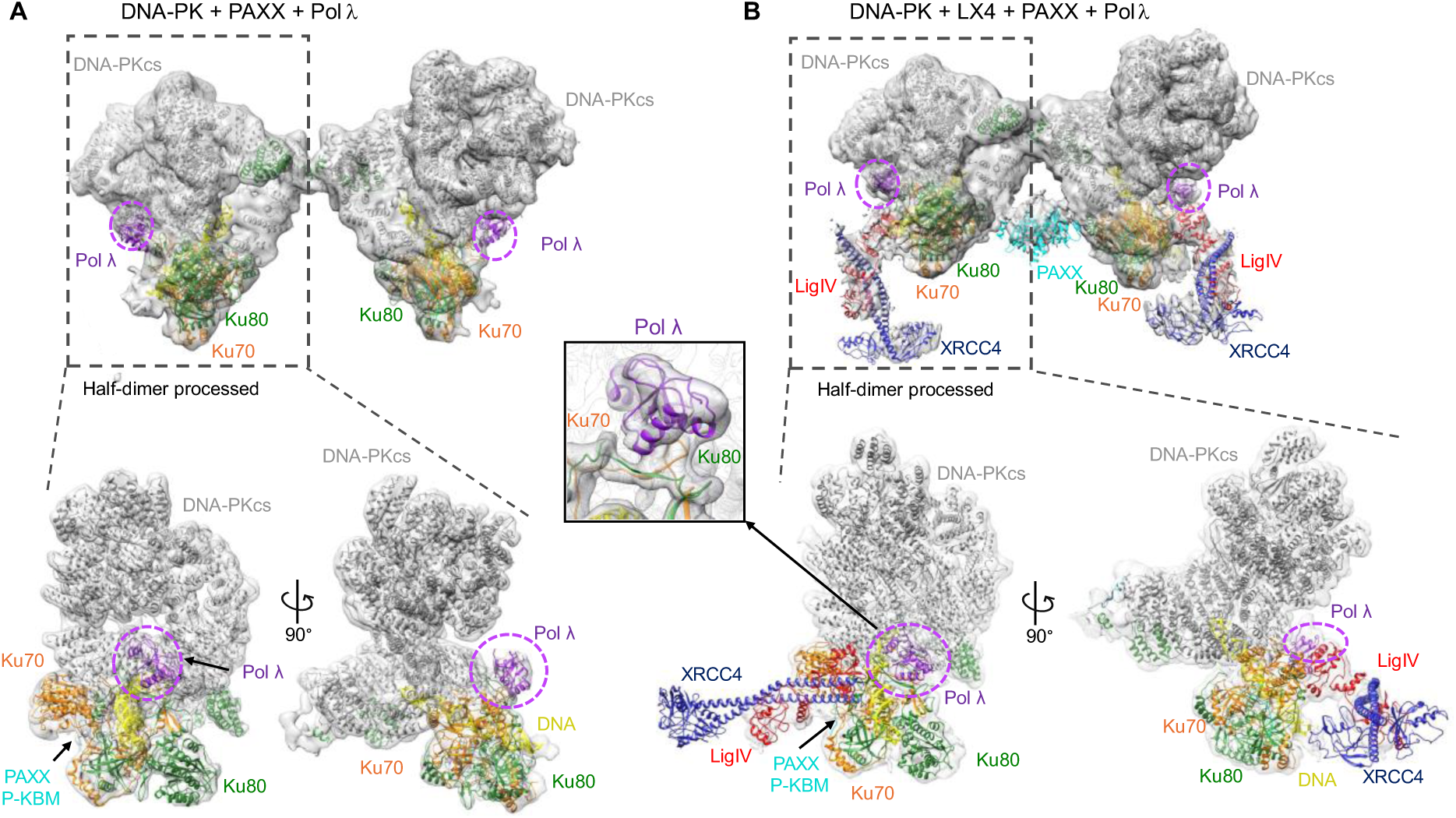
Cryo-EM structures of DNA-PK, PAXX and Pol λ with and without LX4. **A)** DNA-PK + PAXX + Pol λ dimeric map with extra density for Pol λ highlighted in a dashed pink circle, with two orientations of processed half dimer presented below. DNA-PKcs in grey, Ku70 in orange, Ku80 in green, DNA in yellow, PAXX in cyan and Pol λ in purple. **B)** DNA-PK + PAXX + Pol λ + LX4 with extra density for Pol λ highlighted in a dashed pink circle, with two orientations of processed half dimer presented below. DNA-PKcs in grey, Ku70 in orange, Ku80 in green, DNA in yellow, LigIV in red, XRCC4 in dark blue, PAXX in cyan and Pol λ in purple. Inset, zoom in of the Pol λ extra density.

The structure without LX4 bound has an overall resolution of 4.53 Å and the map with LX4 bound has an overall resolution of 4.26 Å **(Figure S2-5)**. Within both of these half-dimer maps, DNA-PK along with the PAXX Ku-binding motif (P-KBM) could be docked **(Figure 1)**. The PAXX P-KBM can be seen interacting with the von Willebrand-like (vWA) domain of Ku70, as has been previously characterised ^2^. Additionally, XRCC4 and BRCT tandem repeats of LigIV can be docked into the map with LX4 bound **(Figure 1)**. As seen in our previous structures there is a central helix in DNA-PKcs which blocks the DNA ends, which is only present when LX4 is engaged. Additional density can be observed at the bridge formed by Ku70 and Ku80 in both maps, into which the NMR structure of the Pol λ BRCT domain (PDB: 2JW5) could be confidently docked **(Figure 1 and S2-5)**. Although the full-length Pol λ was used, only the BRCT domain could be modelled into the cryo-EM map. This suggests that the catalytic domain has no stable interaction with DNA with the LRC in the cryo-EM assemblies determined.

### Molecular basis of the Pol λ BRCT interaction with Ku70/80

The Pol λ BRCT domain is positioned at the interface formed by Ku70 and Ku80, at the periphery of the DNA binding channel formed by the Ku70/80 heterodimer. An interaction between Pol λ and Ku70/80-LigIV-XRCC4 has been previously suggested, with the residues proposed to be involved in the interaction being situated in α1 helix of the BRCT domain ^16^. In agreement with this previous study, our structure shows that Pol λ BRCT domain is positioned to allow helix α1 to dock into a groove formed between Ku70 and Ku80 **(Figure 2D)**. Specifically, the amino acid residues Arg57 and Leu60 of Pol λ BRCT domain appear to mediate the contact with Ku70/80, in agreement with previous data, which found that mutations of these residues prevented Pol λ interaction on DNA with Ku70/80 or Ku70/80-LigIV-XRCC4^16,17^. With regards to Ku70/80, the specific interaction sites encompass residues between 301 – 310 on Ku80 and residues 301 – 308 on Ku70.

**Figure 2.**
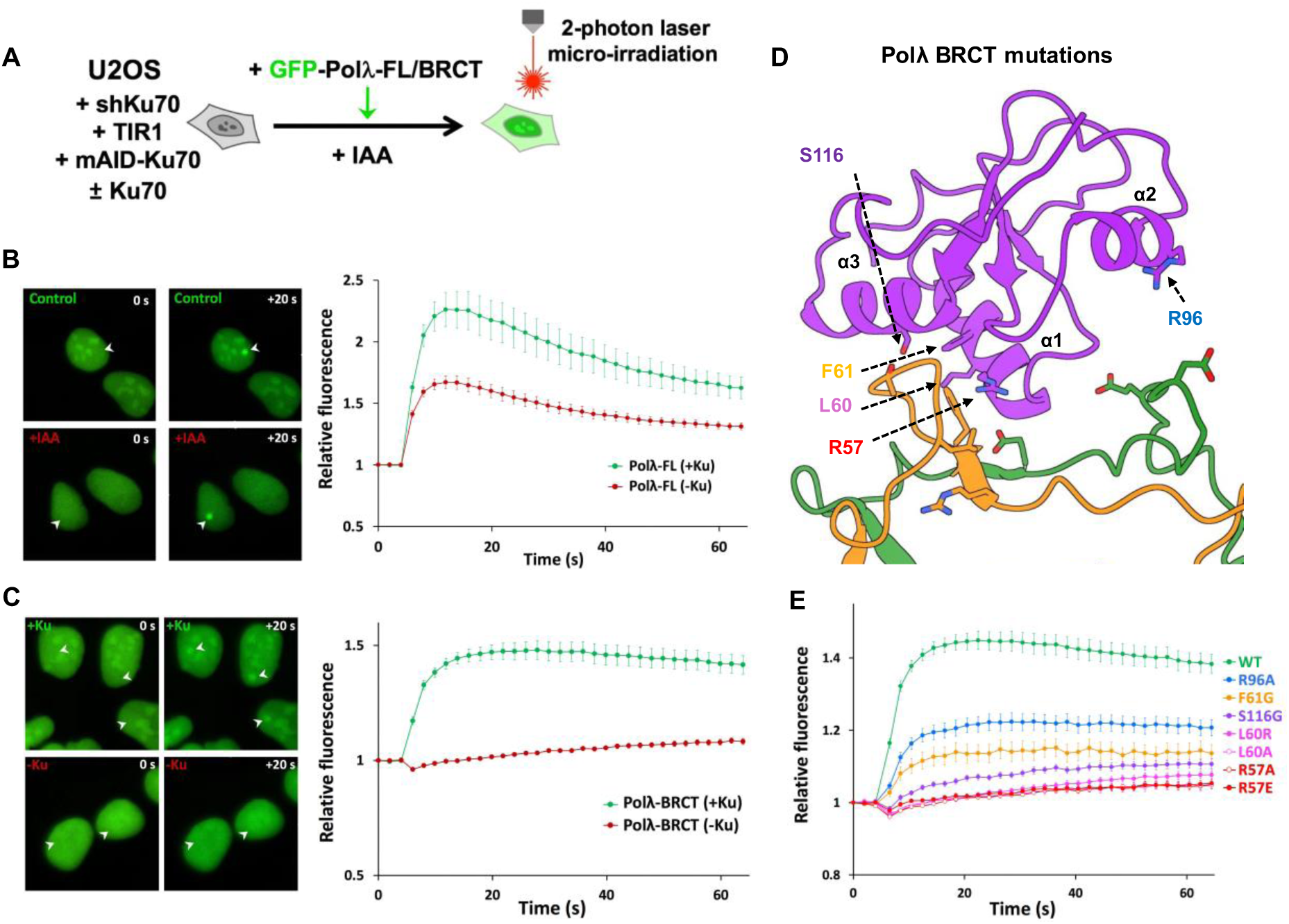
Impact of mutations in the BRCT domain of Pol λ on its recruitment to DSBs by Ku. **A)** Principle of the laser micro-irradiation experiment. U2OS cells engineered for auxin (IAA)-induced Ku70 knockdown, rescued or not with wild-type (WT) Ku70 and expressing either GFP-tagged full-length Pol λ or its BRCT domain, were micro-irradiated with an 800 nm multiphoton laser to generate DSBs in subnuclear areas. **B)** Left panel: representative images before and 20 s after irradiation (irradiated areas are indicated by white arrows) of nuclei from U2OS expressing GFP-tagged full-length Pol λ and depleted or not of Ku70 (±Ku). Right panel: quantification of fluorescence accumulation at laser-induced DNA damage sites. Results are plotted as mean values of at least 20 nuclei ± SEM. **C)** Same as in **B)** with U2OS expressing GFP-tagged Pol λ-BRCT. **D)** Position of mutated residues in the BRCT domain of Pol λ (purple) at the interface with Ku70/Ku80 (orange and green, respectively). **E)** Quantification of fluorescence accumulation at laser-induced DNA damage sites in U2OS cells expressing GFP-tagged WT or mutated BRCT domain of Pol λ. Results are plotted as mean values of at least 20 nuclei ± SEM.

### Mutagenesis of Ku70/80 interaction residues on Pol λ

Based on our structural data, we probed how the Pol λ BRCT interaction with Ku70/80 impacts the response to DNA damage in cells. We first analysed the influence on recruitment of Pol λ at laser-induced DNA damage sites. To establish to what extent Pol λ mobilisation to damaged sites was Ku-dependent, we used in-house engineered U2OS cells that allow circumventing the lethality of Ku loss in human cells as previously described ^2,19^ (**Figure 2 and S6A**). Briefly, endogenous Ku70 expression was first knocked-down via the constitutive expression of an shRNA associated with cell rescue through expression of Ku70 tagged with a mini-auxin inducible degron (mAID). Within a few hours upon addition of auxin (indole-3-acetic acid, IAA), degradation of endogenous Ku70 occurs with concomitant Ku80 depletion due to the known reciprocal stabilization of both Ku subunits. Full-length Pol λ fused to GFP was rapidly and substantially recruited to DNA damage sites in the presence of Ku. This recruitment of the full-length polymerase was only partly Ku-dependent since it substantially persisted under Ku depletion conditions (**Figure 2B, – red line**). This may indicate an ability of Pol λ to interact directly with DNA at sites of damage, or may be related to Ku-independent repair functions of Pol λ outside NHEJ ^20^.

In contrast to data obtained with full-length Pol λ, when the fusion was restricted to the N-terminal Pol λ BCRT domain (**Figure S1A**), its recruitment to micro-irradiated areas was mostly Ku-dependent since it was nearly abolished without Ku (**Figure 2C**). Guided by our structural data, we next probed the impact of specific Pol λ BRCT mutations on the recruitment to DNA damage sites (**Figure 2D)**. As shown in **Figure 2E**, all the mutants tested impaired GFP-Pol λ BRCT accrual at laser-induced DNA damage sites to various extents, with mutations at R57 or L60 positions being the most detrimental.

Notably, while GFP-Pol λ expression was nuclear with a nucleolar enrichment, the latter disappeared upon Ku depletion (**Figure S6B**). The Pol λ BRCT nucleolar enrichment was also strongly reduced with all BRCT mutants tested **(Figure S6C)**. Since it is known that nuclear Ku is enriched in the nucleolus in the absence of DNA damage ^21^, this suggests that GFP-Pol λ localisation to the nucleolus relies on its interaction with Ku, and that this localisation is disrupted by specific mutations within the BRCT domain of Pol λ.

### Mutagenesis of Ku70/80 residues

To identify key residues of Ku70/80 residues at the Pol λ BRCT:Ku interface, we replaced endogenous Ku subunits with mutated forms of either Ku70 or Ku80 as guided by our structural data (**Figure 3A, S7A-C**). We then monitored the impact on Pol λ BRCT recruitment following DNA damage. To ensure that mutants of Ku70/80 did not influence Ku mobilisation at damaged sites *per se*, we monitored simultaneously the recruitment of mCherry-PAXX co-expressed within the same cells, since PAXX recruitment is strictly Ku-dependent ^2^ (**Figure 3B-F**). Mutations on the positions F303, L310 on Ku70 or E292, E304 on Ku80 nearly abolished Pol λ recruitment **(Figure 3D, 3F**). Notably, some subtitutions on these positions (e.g. Ku70 L310R, Ku80 E304A) also affected PAXX recruitment, although at an intermediate extent compared to the full defect observed for Pol λ BRCT **(Figure 3C-E)**. Considering that PAXX binding to Ku70 is far away from the Ku bridge region, this indicates that these substitutions likely compromise the stability of the Ku:DNA interaction in addition to directly affecting the Ku:polymerase interaction. Nevertheless, these data establish that Ku70 F303, L310 and Ku80 E292, E304 are key positions for Ku interaction with Pol λ.

**Figure 3.**
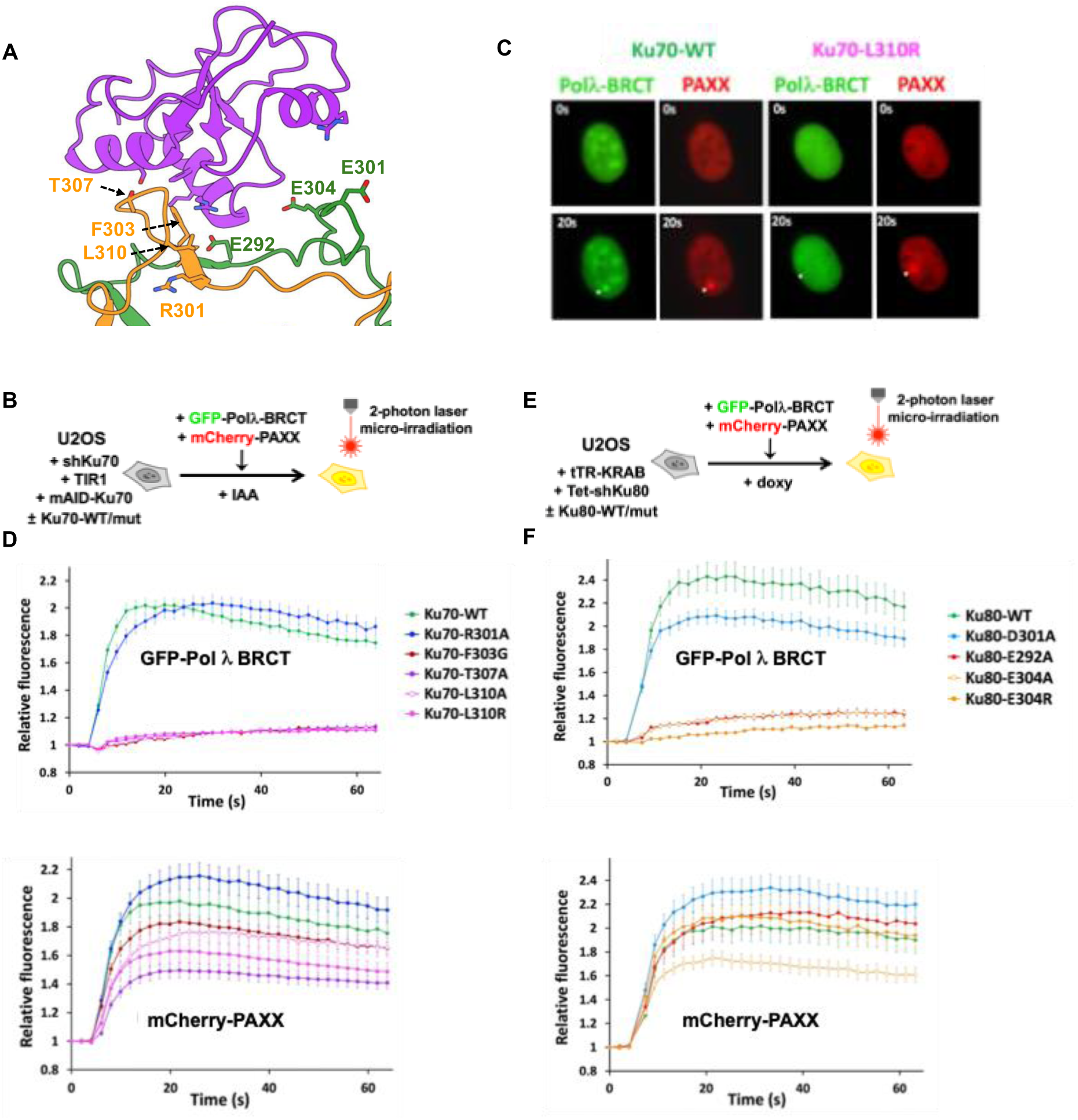
Impact of mutations in the bridge region of Ku70 and Ku80 on the recruitment of the BRCT domain of Pol λ at DSBs. **A**) Position of mutated residues in the bridge domain of Ku70 (orange) or Ku80 (green) at the interface with Pol λ (purple). **B)** Principle of the laser micro-irradiation experiment. U2OS cells engineered for auxin (IAA)-induced Ku70 knockdown, rescued with WT or mutated forms of Ku70 and expressing both the GFP-tagged BRCT domain of Pol λ and the mCherry-tagged PAXX protein, were micro-irradiated and accumulation of each fluorescence was analyzed as in Figure 2. **C**) Representative images before (upper frames) and 20 s after irradiation (lower frames, a white arrow indicates irradiated area) of nuclei from U2OS expressing either WT Ku70 (left) or the L310R mutant (right) **D**) Quantification of fluorescence accumulation at laser-induced DNA damage sites of GFP-Polλ-BRCT (upper chart) and mCherry-PAXX (lower chart) in U2OS cells expressing the indicated Ku70 constructs. Results are plotted as mean values of at least 20 nuclei ± SEM. **E)** Principle of the laser micro-irradiation experiment. U2OS cells engineered for doxycycline (doxy)-induced Ku80 knockdown, rescued with WT or mutants of Ku80 and expressing both the GFP-tagged BRCT domain of Pol λ and the mCherry-tagged PAXX protein, were micro-irradiated and accumulation of each fluorescence was analyzed as in (D). **F**) Quantification of fluorescence accumulation at laser-induced DNA damage sites of GFP-Polλ-BRCT (upper chart) and mCherry-PAXX (lower chart) in U2OS cells expressing the indicated Ku80 constructs. Results are plotted as mean values of at least 20 nuclei ± SEM.

### Gap-filling assay in cells to assess impact of Ku70/80:Pol λ interface mutations

Pol λ has a large spectrum of substrates, accommodating breaks with overhangs (<4 to 12 nt), microhomologies (1-6 nt) and gaps (1-8 nt) ^22^. Based on these features and to assess the impact of mutations at the Ku:Pol λ interface on Pol λ activity during end-joining, we designed a Cas9-targeted plasmid, which once transfected in HEK-293T cells can be used as a gap-filling dependent DNA repair reporter (**Figure 4A**). Briefly, the reporter is designed such that the Cas family member Cpf1 (Cas12a) induces two staggered DSBs in the mCherry cDNA which have two complementary nucleotides at the very end. Gap-filling at the junction restores mCherry expression, while end-filling enables the expression of a downstream EGFP cDNA which lies in a different reading frame. No expression of mCherry or GFP is expected if ends are trimmed or not ligated. Expression of BFP from a co-transfected plasmid accounts for transfection efficiency. Following Cpf1-induced DSB, expression of fluorescent proteins is measured by flow cytometry. We first established that gap-filling dependent mCherry expression relied on cells being transfected by the complete Cpf1/gRNA system (**Figure 4B**), and on NHEJ activity since it was largely inhibited by a DNA-PK inhibitor catalytic activity (NU7441), or upon Ku removal or genetic ablation of the NHEJ factors DNA-PKcs, LigIV, XLF or PAXX involved in break ends tethering and end joining ^2^ (**Figure S7D, E**). Moreover, sequencing the junctions from mCherry positive cells revealed that the expected product from gap-filling repair was formed, further validating our reporter system (**Figure S7F**). Notably, no GFP expression was detected implying that gap-filling is far dominant over end-filling at these staggered DSB (**Figure 4B**). Then, we evaluated the Pol λ-dependency of the gap-filling activity detected. Pol λ was found dominant over Pol µ in gap-filling at three-nucleotide gaps on 5′ overhang substrates ^14,23^. Indeed, Pol λ knock-out (KO) led to a ∼70% reduction of the activity on our substrate that was fully restored upon re-expression of WT Pol λ, while Pol µ KO alone did not impair gap-filling activity (**Figures S7G, H)**. Notably, Pol µ KO in Pol λ KO cells further decreased gap-filling activity, suggesting a small compensation by Pol μ in the absence of Pol λ. Finally, expression of a catalytically dead Pol λ (D427A-D429A, ^24^) (Pol λ dead) further decreased the remaining gap-filling activity in Pol λ KO cells, supporting a dominant negative effect of the Pol λ dead construct on backup gap-filling enzymes, including Pol µ.

**Figure 4.**
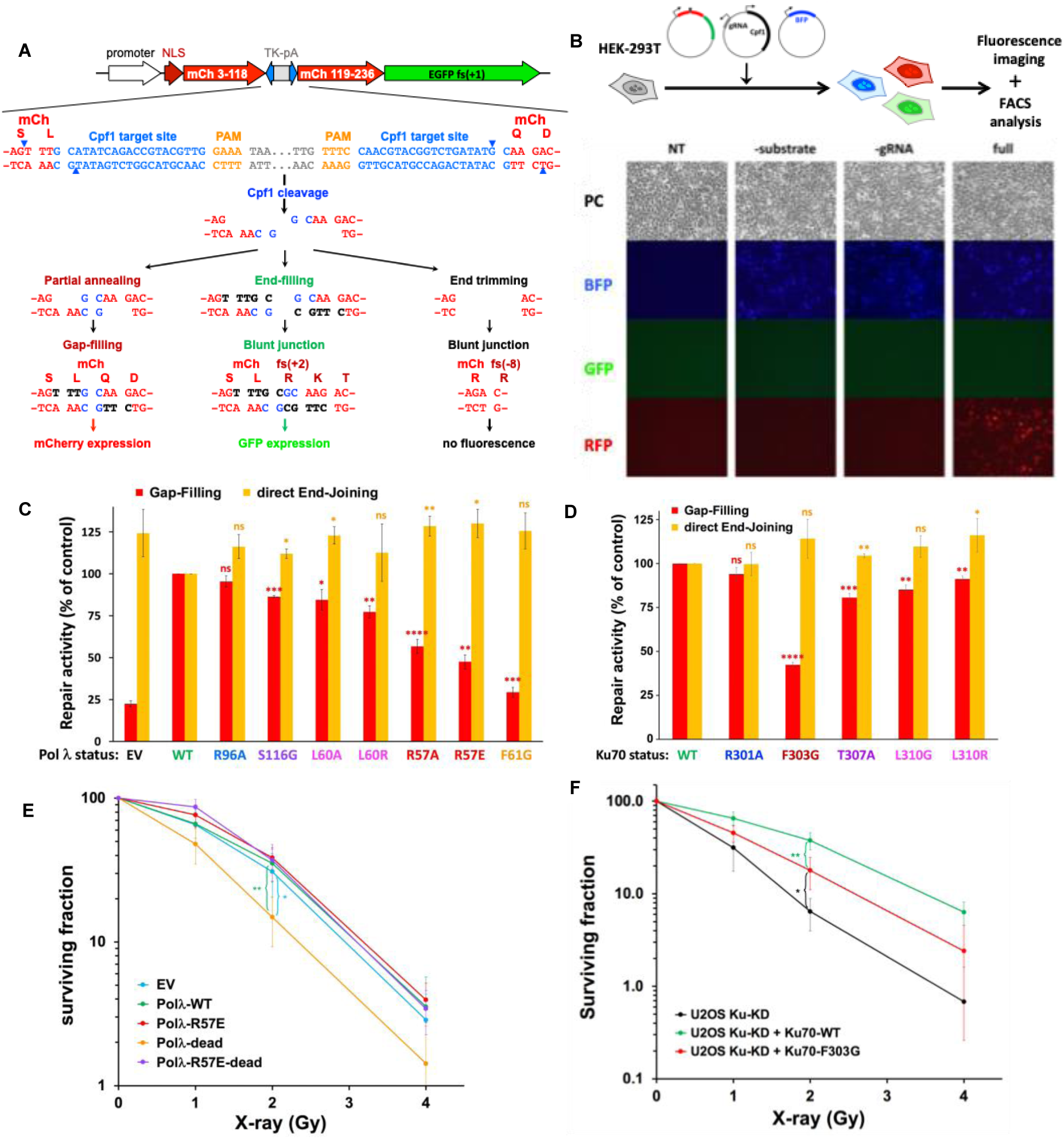
Effect of Pol λ or Ku mutations on gap-filling activity in cells and survival to IR. **A)** Gap-filling reporter substrate. The gap-filling reporter substrate consists of two consecutive frameshifted (+1) coding sequences for mCherry (mCh) and EGFP, respectively. The mCherry coding sequence is interrupted by a cassette containing the HSV-TK polyadenylation sequence (TK-pA) flanked by two inverted copies of a Cpf1 target sequence (blue characters; the PAM GAAA sequence is shown in orange). Following Cpf1-mediated double cleavage, the HSV-TK polyadenylation sequence is deleted and the resulting 5’ overhang DNA ends can be rejoined in different ways. First, the two overhangs can partially anneal to each other by two G:C pairs, leading to single-stranded gaps than can be filled, thus restoring an intact mCherry coding sequence. Second, the two overhangs can undergo end-filling, leading to blunt ends whose joining disrupts the mCherry reading frame (+2 frameshift) but enables the expression of the downstream EGFP coding sequence. Finally, the two overhangs can be trimmed off before end-joining, leading to a -8 frameshift of the mCherry coding sequence and resulting in neither red nor green fluorescence. **B)** Gap-filling assay. Top: the gap-filling reporter assay is performed by transfecting HEK-293T cells with the reporter substrate together with a Cpf1/gRNA-expressing vector to cleave the substrate and a BFP-expressing vector to normalize for transfection efficiency. Fluorescence expression is analyzed 48 h later by flow cytometry. Bottom: representative fluorescence microscopy images of cells non-transfected (NT) or transfected with the full reporter system (full) or omitting the substrate (-substrate) or the gRNA expression vector (-gRNA). PC: phase contrast. **C)** Gap-filling activity (red bars) or direct end-joining activity (orange bars) assessed in parallel in HEK-293T cells knocked-out for Pol λ and complemented with either an empty vector (EV) or expression vectors for WT or different mutants of Pol λ. Results are normalized to the WT condition and plotted as mean values of three to six independent experiments ± SD. P-values from Student’s *t*-test between the WT condition and the considered mutant, for gap-filling and direct end-joining activities, respectively, are as follows: R96A (0.0731; 0.0583), S116G (0.0007; 0.0176), L60A (0.0156; 0.0165), L60R (0.0089; 0.3312), R57A (<0.0001; 0.0025), R57E (0.0021; 0.0252), F61G (0.0005; 0.0528). * P < 0.05, ** P < 0.01, *** P < 0.001, **** P < 0.0001, ns: not significant. **D)** Gap-filling activity (red bars) or direct end-joining activity (orange bars) assessed in parallel in HEK-293T cells knocked-down for Ku70 and rescued with expression vectors for WT or different mutants of Ku70. Results are normalized to the WT condition and plotted as mean values of four experiments ± SD. P-values from Student’s *t*-test between the WT condition and the considered mutant, for gap-filling and direct end-joining activities, respectively, are as follows: R301A (0.0541, 0.9381), F303G (<0.0001; 0.0841), T307A (0.0006, 0.0039), L310G (0.0015; 0.0503), L310R (0.0025; 0.0437). **E)** Cell survival to X-rays of HEK-293T cells knocked-out for POLL and complemented with an empty vector (EV) or expression vectors for WT or the indicated mutants of Pol λ. Y axis is log scale. Results are normalized to the untreated condition and plotted as mean values of four to seven experiments ± SD. P-values were calculated at 2 Gy using unpaired t-test: Polλ-dead versus EV (0.0207); Polλ-dead versus WT (0.0092). **F)** Cell survival to X-rays of U2OS cells knocked-down for Ku70 (Ku-KD) and complemented or not with expression vectors for WT Ku70 or the F303G mutant. Results are mean values of four experiments and plotted as in (E). P-values at 2 Gy: Ku70-F303G versus Ku-KD (0.0195); Ku70-F303G versus Ku-WT (0.0089).

### Characterization of key positions in Pol λ BRCT:Ku interface for Pol λ-dependent gap-filling activity and survival to IR

Using the mainly NHEJ- and Pol λ-dependent gap-filling assay described above, we then analysed the impact of mutations in Pol λ BRCT though transfection of Pol λ KO HEK-293T cells complemented with WT or mutant Pol λ. As a control, the same cells were used to evaluate direct end-joining (EJ) activity at blunt-ended breaks using our dedicated Cas9-based reporter plasmid assay described previously ^19^. As shown in **Figure 4C**, except for R96A, all other mutations tested in Pol λ BRCT reduced gap-filling efficiency at staggered DSB. The effect of F61G mutation is inconclusive since it lowered the protein expression **(Figure S8A)**. Notably, the extent of defect observed in gap-filling activity of full-length mutant Pol λ correlates with that observed in recruitment of the corresponding mutant GFP-BRCT at laser sites (**Figure 2E**), again with R57 and L60 positions being the most crucial. Since no detectable repair defect was found at blunt-ended DSB, a readout for direct EJ, this indicates that outside its catalytic function at defined breaks Pol λ is unlikely to fulfill a general function in the overall NHEJ complex assembly and/or stability. Using the same assays, we then evaluated the impact of mutations of Ku70 in the Pol λ:Ku interface by using HEK-293T cells expressing mAID-Ku70 fusion and transduced with WT or mutated Ku70 forms (**Figure S8B**). Upon auxin addition, we analysed in parallel the impact of individual or combined mutations in Ku70 on gap-filling at staggered DSB and on direct EJ at blunt DSB (**Figure 4D**). Compared to the slight reduction in gap-filling observed with Ku70 mutants at T307 and L310 positions, F303G mutation reduced gap-filling by 60% without impairing repair at blunt-ended breaks, indicating that Ku70 F303 is a crucial position for Ku interaction with Pol λ BRCT but not for Ku interaction with DNA. Then, assessing cell radiosensitivity, we observed that loss of Pol λ marginally reduced cell resistance to ionizing radiation (IR) that was much more decreased upon expression of the Pol λ dead construct (**Figure 4E**), in agreement with gap-filling activity assessed in parallel (**Figure S7H**). Notably, the R57E mutation that impaired Pol λ recruitment to DSB released the sensitivity associated with expression of the Pol λ dead construct when both mutations were combined (**Figure 4E**) and similarly enhanced gap-filling activity up to the level observed in Pol λ KO cells (**Figure S8C**). Finally, since we observed that the loss of cell viability consecutive to Ku removal is reversible within a 32 h time window, we assessed the consequence of impairing Ku:Pol λ interaction on cell survival to IR (**Figure S8D and Figure 4F**). We showed that the Ku70 F303G mutant that preserved direct EJ but impaired gap-filling did not fully complement the radiosensitivity observed upon Ku removal. This supports that Ku interaction with gap-filling polymerases promotes cell survival to IR through positioning them at the break ends (**Figure 4F**).

## Discussion

We determined the cryo-EM structures of Pol λ in complex with the Ku70/80 heterodimer bound to DNA whilst engaged with the DNA-PK holoenzyme. These structures show a clearly defined interaction between the BRCT domain of Pol λ and the interface between Ku70/80. We assessed the functionality of this interaction by generating specific site directed mutants on either Pol λ BRCT or Ku70/80. These mutants were used in two orthogonal assays: live protein recruitment at nuclear laser sites and gap-filling activity in cells transfected with an original reporter assay based on Cpf1-generated partially complementary DSB ends. These data define the molecular basis and essentiality of the BRCT domain for recruitment of Pol λ within the NHEJ complex. The data also establishes key positions on Ku70/80 and Pol λ-BRCT that mediate their interaction, and for the first time position the interaction region at the external face of the Ku70/80 dimer interface.

From our cryo-EM structure of the Ku80 mediated DNA-PK dimer bound to LX4, the LigIV tandem BRCT1 domain can be seen occupying the previously described site on Ku70/80 ^6,7^ that is distinct from the Pol λ BRCT interaction site **(Figures 1 and 5**). The LigIV BRCT1 sits in the groove formed by the Ku70/80 dimer interface, whereas Pol λ BRCT is located at an adjacent site. From our structural data, it is apparent that the two proteins have distinct sites of contact with Ku70/80, and there is no evidence that Pol λ and LigIV form direct interactions with each other. Thus, the previously reported observation that specific Pol λ mutations prevent complex formation with Ku70/80-LigIV-XRCC4 are likely due to disruption of the Pol λ:Ku70/80 interaction specifically, as has previously been suggested ^16^. Interestingly, when we collected cryo-EM data of Ku70/80 alone (without DNA-PKcs) with Pol λ, we did not observe any density for the BRCT domain of Pol λ engaged with Ku70/80. Therefore, it is possible that the Pol λ interaction with Ku70/80 is stabilised within the DNA-PK holoenzyme complex. Our cryo-EM structures indicate a possible weak interaction between the BRCT domain of Pol λ and DNA-PKcs. Although the resolution is low, we would predict that Pol λ (residues 128-130) may interact with residues Ser187 and/or Glu188 on DNA-PKcs.

**Figure 5:**
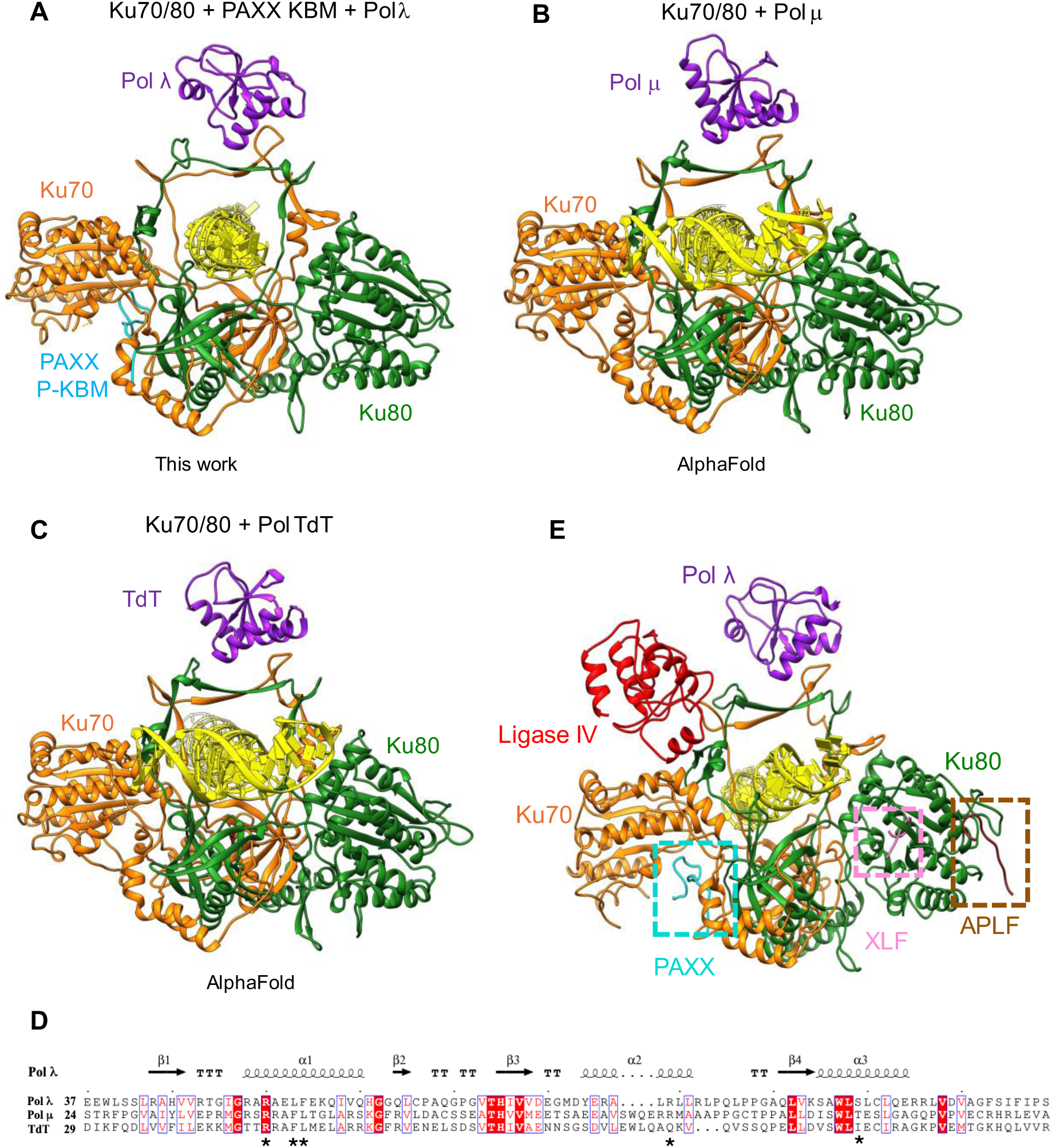
Comparison of the binding of BRCT domains from NHEJ polymerases X (λ, μ and TdT) to Ku70/80 and scheme of Ku as a structural hub. **A)** Experimentally determined structure from this work showing Pol λ binding to Ku70/80, DNA and PAXX. **B)** AlphaFold 3 prediction of Pol μ binding to Ku70/80 and DNA. **C)** AlphaFold 3 prediction of Pol TdT binding to Ku70/80 and DNA. Ku70 is in orange, Ku80 in green, DNA in yellow, PAXX P-KBM in cyan and Polymerase BRCTs in purple. **D)** Sequence alignment of the BRCT domain of Pol X family members generated by ESPript ^30^. Structural elements from the Pol λ cryo-EM structures are shown above the alignment. Residues mutated in this study are indicated with *. **E)** Ku70/80 (orange/green) as a structural hub showing the binding of Pol λ (purple), Ligase IV (red), PAXX (cyan), XLF (pink) and APLF (brown).

Notably, we found that recruitment of the N-terminal Pol λ BRCT domain to micro-irradiated areas was essentially Ku70/80-dependent in contrast with data obtained with the full-length Pol λ. This suggests that Pol λ recruitment to sites of DNA damage may rely on interactions outside of its BRCT domain. This may reflect additional roles of Pol λ outside of classical NHEJ such as in Base Excision Repair where it is proposed to interact with DNA glycosylases involved in the repair of alkylated or oxidized bases ^20^. As previously reported, we found that Pol λ deficient cell lines showed modest, if any, sensitivity to IR, whereas cells deficient in both λ and µ polymerases were clearly IR sensitive ^14,25,26^. The expression of a catalytically inactive form of Pol λ concurrently inhibited gap-filling activity in cells (this work) and negatively impacted on cell survival to IR as previously reported ^24^, to extents higher than the sole deletion of Pol λ. This suggests that Pol λ dead prevents the rescue of the Pol λ defect by Pol µ, likely through occupying and occluding a common interaction site on Ku70/80, and thereby sustaining a dominant negative effect. Also, we showed that Ku70 F303G mutant cells exhibit sensitivity to IR equivalent to that of a double Pol λ-Pol μ KO cells, suggesting again that this mutation impairs Ku70/80 interaction with both Pol X proteins. Indeed, AlphaFold prediction of the structures formed between Ku70/80 with DNA and the BRCT domains of Pol μ and TdT indicates the use of similar protein:protein interfaces (**Figure 5B, C**). Also, the broad composition and organization of the Pol λ BRCT domain is conserved amongst Pol X members, supporting a conserved mode of interaction with Ku70/80 **(Figure 5D)**. Together with our present cellular data, this allows us to propose a unified model for the interaction of the three BRCT-bearing Pol X with Ku70/80 on the same site allowing their respective recruitment to the NHEJ complex.

The Ku70/80 interaction with Pol X BRCT domain reported here adds to the fascinating list of Ku70/80 sites that contact NHEJ factors. These include Ku80 sites for A-KBM bearing proteins (APLF, WRN, MRI), XLF via its X-KBM ^27^ and DNA-PKcs ^8^, Ku70 sites for PAXX via its P-KBM ^2^ and LigIV via its BRCT1 ^7,8^, and combined sites in Ku80/Ku70 for inositol hexaphosphate ^19^ and now for Pol X BRCTs (B-KBM). The structures reported here of the Ku70/80:Pol λ interaction represents a paradigm for Pol X mode of binding to Ku70/80 and reinforces the notion that Ku70/80 acts as a central structural hub in the NHEJ mechanism ^28^. Like other Ku70/80 partners whose KBM motifs are at the N or C-terminus of the protein, Pol X members are anchored by an N-terminal BRCT domain separated by a flexible region from the catalytic portion of the protein, allowing free movement for the correct positioning at the site of the DSB.

The structures of the LRC presented here indicate that the Pol X:Ku70/80 interaction occurs early during the NHEJ process, at a stage where DNA-PKcs is still present and at which the DNA ends are likely not yet accessible to processing enzymes. Moreover, since our cryo-EM structures were obtained using a blunt-ended DNA substrate, this suggests that it is not the nature of the DNA ends that dictates Pol X engagement into the NHEJ process but rather its intrinsic affinity for the initial DNA-PK complex. These structures also illustrate that the evolution of distinct and independent binding sites on the Ku70/80 heterodimer enables the concurrent anchoring of NHEJ proteins (PAXX/XLF, DNA-PKcs, Pol X, XRCC4/LigIV) dedicated to specific activities involved in the repair reaction (synapsis, kinase, polymerase, ligase, respectively) **(Figure 5E)**. Moreover, the dimeric nature of the LRC allows for a scenario where two different members of Pol X family could be engaged within the same dimeric complex simultaneously. Although our DSB repair assay in cells could theoretically report both gap-filling and end-filling activities, we found that gap-filling is far dominant. This suggests that, when possible, end-annealing on minimal homology is preferred, most likely because it contributes to the stability of the synaptic complex and ultimately to the fidelity of DNA repair. The two-nucleotide overhang substrate used here supports a tight coupling between end annealing and gap filling, as demonstrated elegantly for NHEJ mediated repair in Xenopus egg extracts ^23^. Within the NHEJ LRC, this coupling is ensured by the concurrent binding of ends, and resynthesis factors by Ku70/80 on both sides of the DSB. Also, the coexistence of LigIV and processing enzymes like Pol X in the synaptic complex at break ends ensures that ligation can proceed as soon a DNA ends are ligatable, limiting unnecessary DNA sequence alteration ^23^. This illustrates that the unique capability of Ku70/80 for simultaneous multivalent interaction is crucial for the high adaptability of the NHEJ process ^29^.

## Acknowledgements

We thank Dr Christos Savva, Dr Emma Hesketh, Dr Claudia Lancey and Dr TJ Ragan from the Midlands Regional Cryo-EM facility for help with grid preparation, screening, data collections and processing support. P.F., N.B., J.C., S.B. V.R, J.B.C and P.C. were supported by the French National Research Agency (ANR-20-CE11-0026). This work was supported by the Fondation ARC (J.C.). P.C. is a scientist from INSERM. We acknowledge the imaging facility TRI, member of the national infrastructure France-BioImaging infrastructure supported by the French National Research Agency (ANR-10-INBS-04). JB.C and V.R thank the I2BC Protex platform supported by French Infrastructure for Integrated Structural Biology (FRISBI) ANR-10-INBS-0005. J.C and V.R were supported by ANR-21-CE12-0019-01, ANR-22-CE12-0037 and ANR 23-CE11-0033.

## Author contributions

A.K.C. directed the study. A.K.C., P.C. and P.F. led the experimental design. H.A., S.Z., A.K.C., S.B., P.F. and P.C. prepared the manuscript. H.A. and S.Z. collected the cryo-EM data and modelled and analysed the structures. C.H. helped with structural analysis and edited manuscript. G.M. Purified proteins and collected initial cryo-EM data. P.F. designed and built the cellular tools. P.F. and N.B. designed, validated and carried out the in vivo repair experiments. P.F. and J.C. performed the multiphoton laser micro-irradiation experiments. A.K.C., S.B. and P.C. acquired funding. S.W.H and D.Y.C advised with cryo-EM data collection set-up and processing. S.W.H helped with structural analysis and manuscript preparation. V.R and J.B.C provided insect cells pellets for human NHEJ factors.

## Data and materials availability

All structural data presented are publicly available. Cryo-EM structures and maps are deposited at the PDB and EMDB with accession codes as follows:

## Materials and Methods

### Purification of DNA-PKcs and Ku70/80

DNA-PKcs and full-length His-tagged Ku70/80 were expressed and purified as previously described ^6^.

### Expression and purification of full-length XLF and LX4

A construct containing full-length 10xHis-tagged XLF and LX4 (LigIV and XRCC4) were expressed in insect cells and purified as previously reported ^2^.

#### DNA annealing

Biotinylated Y-shaped 42-55 bp dsDNA were synthesised and annealed as described previously ^6^. DNA sequences used for annealing can be found below.

Y-shaped DNA Forward

Biotin-

CGCGCCCAGCTTTCCCAGCTAATAAACTAAAAACTATTATTATGGCCGCACGCGT

Y-shaped DNA Reverse ACGCGTGCGGCCATAATAATAGTTTTTAGTTTATTGGGCGCG

### Cryo-EM sample preparation of DNA-PK, LX4, PAXX and Polymerase λ structure

Proteins were concentrated using a centrifugal filter (Amicon) with a 30 kDa cut-off and buffer exchanged into 20 mM HEPES, pH 7.6, 200 mM NaCl, 0.5 mM EDTA, 2 mM MgCl_2_, 5 mM DTT. Purified Ku70/80 full-length was then first mixed with Y-shaped 42-55 bp DNA before being mixed with purified DNA-PKcs, LX4, PAXX and Pol λ in respectively a 1:1:2:2:2:6 (DNA:DNA-PKcs:Ku:LX4:PAXX:Polλ) ratio.

### Cryo-EM grid preparation

Aliquots of 3 μl of ∼2.5 mg/ml of the NHEJ complex (based on DNA-PKcs concentration) were mixed with 8 mM CHAPSO to eliminate particle orientation bias (final concentration; Sigma) before being applied to Holey Carbon grids (Quantifoil Cu R1.2/1.3, 300 mesh), glow discharged for 60 s at current of 25 mA in PELCO Easiglow (Ted Pella, Inc.). The grids were then blotted with filter paper once to remove any excess sample, and plunge-frozen in liquid ethane using a FEI Vitrobot Mark IV (Thermo Fisher Scientific) at 4 °C and 95 % humidity.

### Cryo-EM data acquisition

The data was collected on a Titan Krios equipped with a Gatan K3 direct electron counting detector at the University of Leicester. All data collection parameters are given in **Table S1**.

### Cryo-EM Image processing

The classification process for the two datasets is summarised schematically in **Figure S2**. The final reconstructions obtained had overall resolutions **(Table S1**), which were calculated by Fourier shell correlation at 0.143 cut-off.

### Cryo-EM structure refinement and model building

The model of the DNA-PK monomer (PDB:7NFE) was used as an initial template and rigid-body fitted into the cryo-EM density in UCSF chimera ^31^ and manually adjusted and rebuilt in Coot ^32^. LX4 was either removed or manually adjusted depending on whether density was present and PAXX was docked within the central density. The BRCT domain of Polymerase λ was then manually docked into the density (PDB: 2JW5) in chimera. Models and their corresponding density maps were run through Namdinator ^33^ before being refined using Phenix real-space refinement ^34^.

### Cell lines, cell culture and cell engineering

U2OS cells (human osteosarcoma cell line from ECACC, Salisbury, UK) and HEK-293T human embryonic cells, were grown in DMEM (Eurobio, France) supplemented with 10% fetal calf serum (Eurobio, France), 125 U/ml penicillin, and 125 μg/ml streptomycin. Cells were maintained at 37°C in a 5% CO2 humidified incubator.

HEK-293T cells knocked-out for POLL (DNA Polymerase λ), POLM (DNA Polymerase μ), PRKDC (DNA-PKcs, Addgene Plasmid#220493 ^35^, LigIV, XLF and PAXX genes were obtained following cell transfection with the pCAG-eCas9-GFP-U6 vector expressing the corresponding guide RNA (see below) using jetPEI (Polyplus) as a transfection reagent. Following cell sorting, individual clones were isolated and checked by western blot.

The generation of U2OS cells expressing an inducible shRNA against Ku80 and the generation of U2OS and HEK-293T cells expressing a mini-auxin-inducible degron-tagged Ku70 protein (mAID-Ku70) in place of the endogenous Ku70 protein have been previously described ^19,36^. Production of lentiviral particles in HEK-293T cells and transduction of U2OS and HEK-293T cells were performed as previously described ^37^.

### Western-blot and antibodies

Cell pellets were washed with phosphate-buffered saline (PBS) and resuspended in lysis buffer (50 mM Hepes-KOH, pH 7.5, 450 mM NaCl, 1 mM EDTA, 1% Triton X-100) supplemented with Halt protease inhibitor cocktail (ThermoFisher Scientific). Cells were lysed by four freeze/thaw cycles in liquid nitrogen and 37°C water bath. Lysates were cleared by centrifugation and protein concentrations were determined using the Bradford assay (Bio-Rad, Hercules, CA). Equal amounts of proteins were mixed with concentrated loading sample buffer to 1X final concentration (50 mM Tris.HCl pH 6.8, 10 % glycerol, 1 % SDS, 300 mM 2-mercaptoethanol, 0.01 % bromophenol blue), heat-denatured, separated by SDS-PAGE on Miniprotean TGX stain-free 4-15 % gradient gels (Bio-Rad, Hercules, CA) and blotted onto Protran 0.45 µm nitrocellulose membranes (GE Healthcare). Membranes were blocked for 60 min with 5% non-fat dry milk in PBS, 0.1% Tween-20 (Sigma-Aldrich) (PBS-T buffer), incubated as necessary with primary antibody diluted in PBS-T containing 1% bovine serum albumin (immunoglobulin- and lipid-free fraction V; Sigma-Aldrich) and washed 3 times with PBS-T. Membranes were incubated for 1 h with HRP-conjugated secondary antibodies (Jackson Immunoresearch Laboratories) in PBS-T and washed five times with PBS-T. Immuno-blots were visualized by enhanced chemiluminescence (Western Lightning Plus-ECL; Perkin Elmer) and autoradiography. Primary antibodies used: mouse monoclonal antibodies anti-DNA-PKcs (clone 18.2; Thermo Fisher Scientific), anti-Ku80 (clone 111), anti-Ku70 (clone N3H10), anti-Pol lambda (clone E11, Santa Cruz), anti-beta-Actin (clone C4, Santa Cruz); rabbit monoclonal antibodies anti-LigIV (A11432, Abclonal), anti-Pol mu (EPR10470(B), Abcam); rabbit polyclonal antibodies anti-XLF (A199957, Abclonal), anti-PAXX (NBP1-94172, Novus).

### Ionizing irradiation and cell survival analysis

Two to six thousand U2OS or HEK-393T cells per well were seeded in duplicate in six-well plates. Cells were exposed 24 h later to various doses of X-ray using a Faxitron RX-650 device (130 kV, 5 mA, dose rate 0.3 Gy per min). Six to seven days later, cells were fixed with 7% trichloroacetic acid for 1 h at 4°C. Fixed cells were extensively washed with water and plates were air dried before staining for 15 min with crystal violet (0.1% aqueous solution). Stained cells were further extensively washed with water and plates were air dried. Staining was dissolved with 10% acetic acid solution and absorption was measured at 570 nm (Ultrospec-3000 spectrophotometer, Pharmacia Biotech). Results were plotted as mean values of >3 independent experiments ± s.d. using Microsoft Excel software.

### Multiphoton laser micro-irradiation

Live cell microscopy and multiphoton laser micro-irradiation were conducted as previously described ^27^.

### *In vivo* DNA end-joining assays

To assess gap-filling activity, HEK-293T cells were seeded to 20-40% confluence in 6-well plates and transfected 24 h later with a mix of Cpf1-targeted gap-filling reporter substrate, Cpf1/gRNA expressing vector and mTagBFP2 expressing plasmid as an internal control. Cells were trypsinized 2 days post-transfection, washed with PBS and analyzed by flow cytometry on a Fortessa X-20 cell analyzer (BD Biosciences). The integrated red fluorescence signal accounting for gap-filling-mediated repair events (% positive cells x mean fluorescence) was normalized to that of transfection efficiency (BFP). For the repair junction analysis, a variant of the substrate vector was generated in which a sequence encoding HygroR-T2A was inserted in frame, upstream of the disrupted mCherry sequence. This plasmid was stably transfected into HEK-293T cells and, after selection with hygromycin, positive clones were isolated. Following further transient transfection with the Cpf1/gRNA expressing vector, red fluorescent and non-fluorescent clones were isolated. Sequence junctions were analyzed by genomic DNA extraction (SV Genomic DNA Purification System, Promega), PCR amplification with primers mCh-Xba-F and mCh-Xho-R and sequencing (Eurofins Genomics; Ebersberg, Germany). Direct end-joining activity was assessed as described previously with a dedicated reporter substrate in which blunt-ended DSBs were generated with appropriate Cas9/gRNA co-expression ^19^.

### Plasmids and DNA manipulations

Cas9 and gRNA expressing vectors for gene knockout were generated by inserting the pre-annealed gRNA-F and gRNA-R oligonucleotides of the corresponding targeted gene into the BbsI restriction sites of pCAG-eCas9-GFP-U6-gRNA plasmid (a gift from Jizhong Zou, Addgene plasmid # 79145 ; http://n2t.net/addgene:79145 ; RRID:Addgene_79145). See below the list of oligonucleotides.

The previously described pLV3 lentiviral vector ^27^ was modified as follows to allow insertion of Pol λ cDNA downstream a puromycin resistance gene and a sequence encoding the T2A ribosomal skipping peptide: first, a T2A cassette (pre-annealed oligonucleotides kpn2-T2A-Mlu-F and kpn2-T2A-Mlu-R) was inserted into the Kpn2I and MluI restriction sites of the pLV3 plasmid; second, a PCR-amplified fragment with a Puro-resistance cDNA sequence (PCR reaction with primers HF-Puro-F and HF-Puro-R on a synthetic DNA molecule as a template) was added by Hot-Fusion ^38^ at the Kpn2I site, resulting in the pLV3-Puro-T2A plasmid. A PCR-amplified human Pol λ cDNA fragment (primers PolL-Mlu-F and PolL-Bcu-R) was then inserted between the MluI and BcuI restriction sites of pLV3-Puro-T2A. Expression vectors for mutant forms of Pol λ were obtained in a similar manner following an additional step of overlap extension PCR mutagenesis with the corresponding PolL-mut-F and PolL-mut-R oligonucleotides as mutated inner primers (see below the list of primers).

The expression vector for GFP-tagged Pol λ constructs (full-length protein or BRCT domain) was obtained by replacing the FLAG-Ku70 cDNA from the previously described pLV3-GFP-FLAG-Ku70 vector ^2^ by a linker cassette (pre-annealed Kpn2-AX-Mlu-F and Kpn2-AX-Mlu-R oligonucleotides) between the Kpn2I and MluI sites. The resulting pLV3-GFP plasmid was then used to insert between the MluI and BcuI sites the PCR-amplified human Pol λ cDNA fragment described above. Expression vectors for GFP-tagged WT and mutant Pol λ BRCT domain (aminoacids 1-136) were obtained in a similar manner following a PCR amplification of the corresponding cDNAs with PolL-Mlu-F and PolL-BRCT-Bcu-R primers and full-length Pol λ expression vectors as templates.

The lentiviral vector allowing expression of mCherry-tagged human PAXX was obtained, first, by inserting between Kpn2I and MluI restriction sites of pLV3 the coding sequence of mCherry, following PCR amplification from the pmCherry-NLS plasmid (a gift from Martin Offterdinger; Addgene plasmid # 39319 ; http://n2t.net/addgene:39319 ; RRID:Addgene_39319) with primers mCh-Kpn2-F and mCh-Mlu-R, second, by further inserting between MluI and BcuI restriction sites the coding sequence of PAXX, following PCR amplification from the previously described pLV3-GFP-PAXX plasmid ^2^ with primers PAXX-Mlu-F and PAXX-Bcu-R.

Ku70 lentiviral expression vector was generated by PCR amplification of human Ku70 cDNA with Ku70-Kpn2-F and Ku70-Bcu-R primers and subsequent insertion between Kpn2I and BcuI restrictions sites of the pLV3 vector. Expression vectors for mutant forms of Ku70 were obtained in a similar manner following an additional step of overlap extension PCR mutagenesis with the corresponding Ku70-mut-F and Ku70-mut-R oligonucleotides as mutated inner primers (see below the list of primers).

Ku80 lentiviral expression vector was already described ^19^. Expression vectors for mutant forms of Ku80 were generated following an additional step of overlap extension PCR mutagenesis with the corresponding Ku80-mut-F and Ku80-mut-R oligonucleotides as mutated inner primers (see below the list of primers).

The gap-filling reporter substrate was assembled into the pEGFP-N1 vector (Clontech) following sequential insertion of various PCR products, oligonucleotide linkers and synthetic DNA fragments (see **Figure 4A** for a detailed description). The Cpf1 and guide RNA co-expression plasmid used to cleave the reporter substrate was generated by inserting the pre-annealed gRNA-GF-F and gRNA-GF-R oligonucleotides into the Esp3I restriction sites of pTE4398 (a gift from Ervin Welker ; Addgene plasmid # 74042 ; http://n2t.net/addgene:74042; RRID:Addgene_74042).

The pNLS-mTagBFP2 plasmid used as an internal control of transfection efficiency was obtained by PCR amplification of the 2xNLS-mTagBFP2 coding sequence with mTagBFP-Acc65-F and mTagBFP-Mlu-R primers on the pHAGE-TO-nls-st1dCas9-3nls-3XTagBFP2 plasmid template (a gift from Thoru Pederson ; Addgene plasmid # 64512; http://n2t.net/addgene:64512 ; RRID:Addgene_64512). The resulting PCR fragment was inserted into the pEGFP-N1 (Clontech) vector backbone after modification of the multiple cloning site and removal of the GFP coding sequence.

All oligonucleotides were purchased from Eurofins Genomics (Ebersberg, Germany). Restriction and modifying enzymes (Phusion and T4 DNA Ligase) were from ThermoFisher Scientific (Illkirch, France). All constructs were checked by sequencing (Eurofins Genomics).

## Oligonucleotides (DNA linkers and PCR primers)

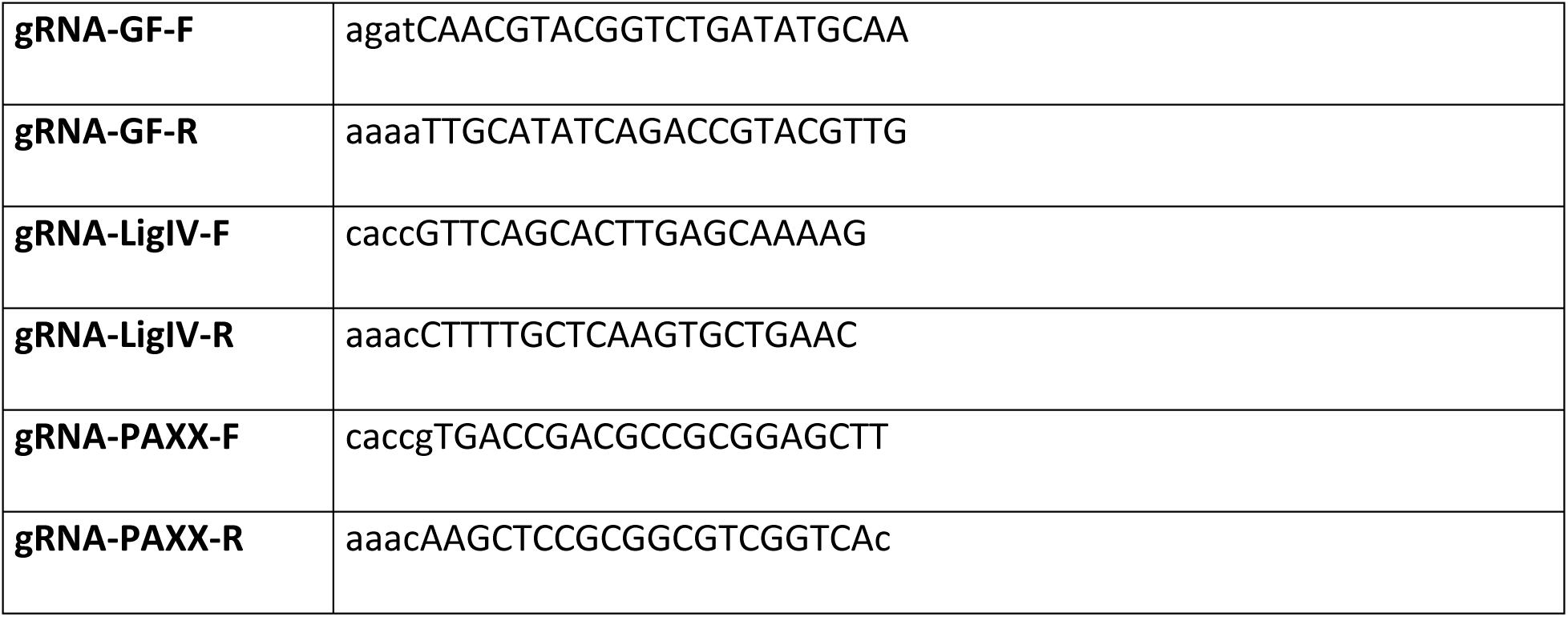
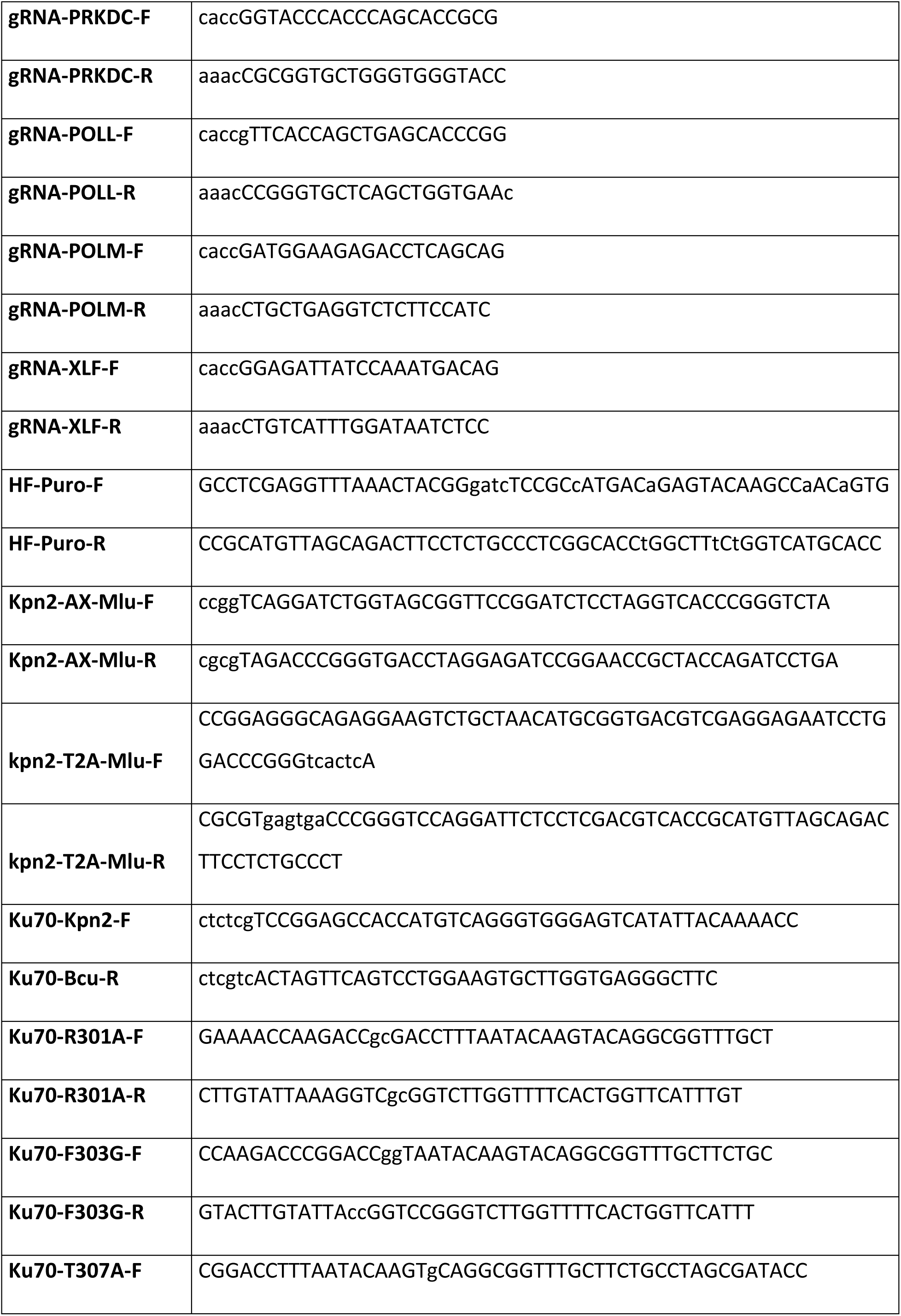
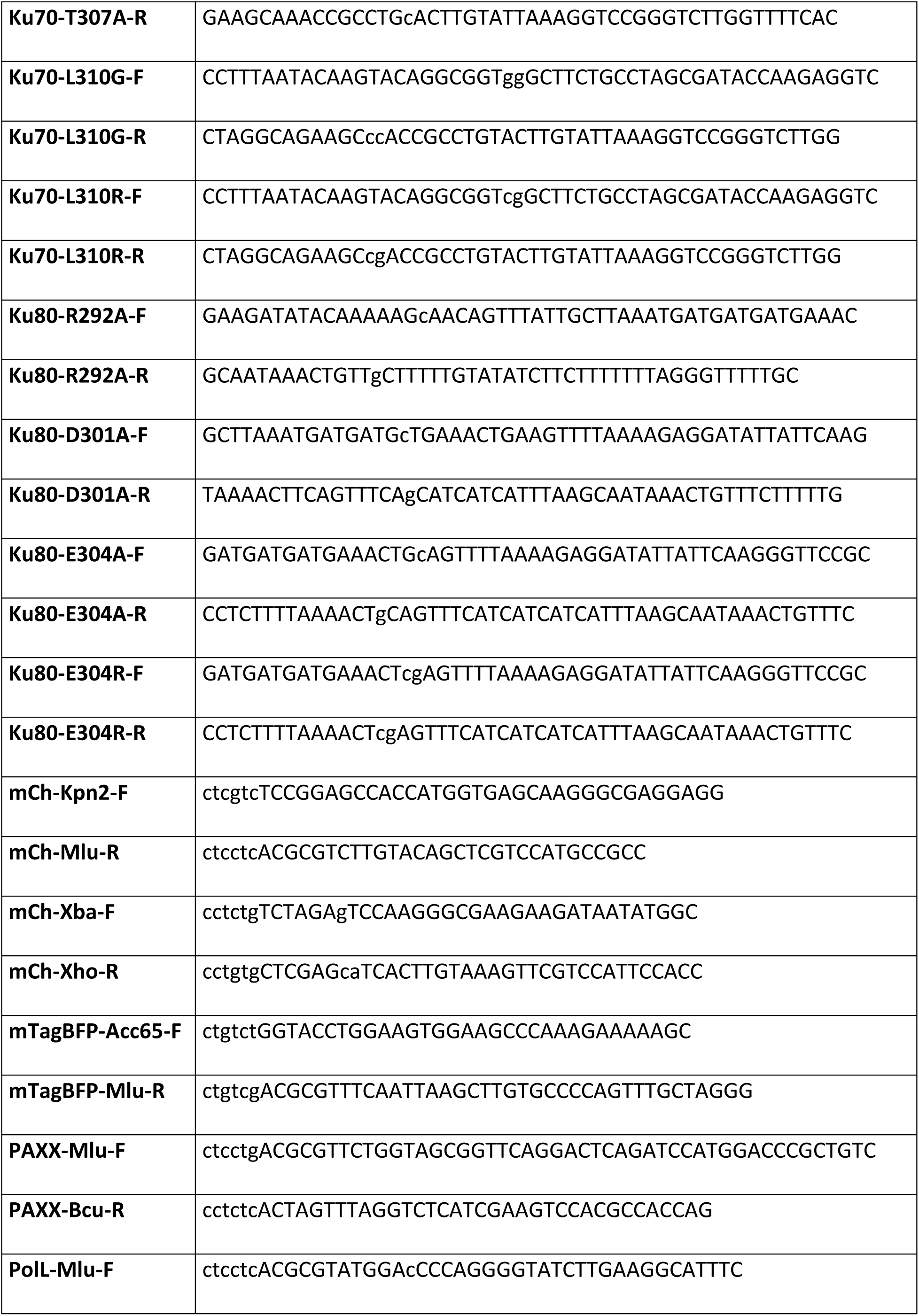
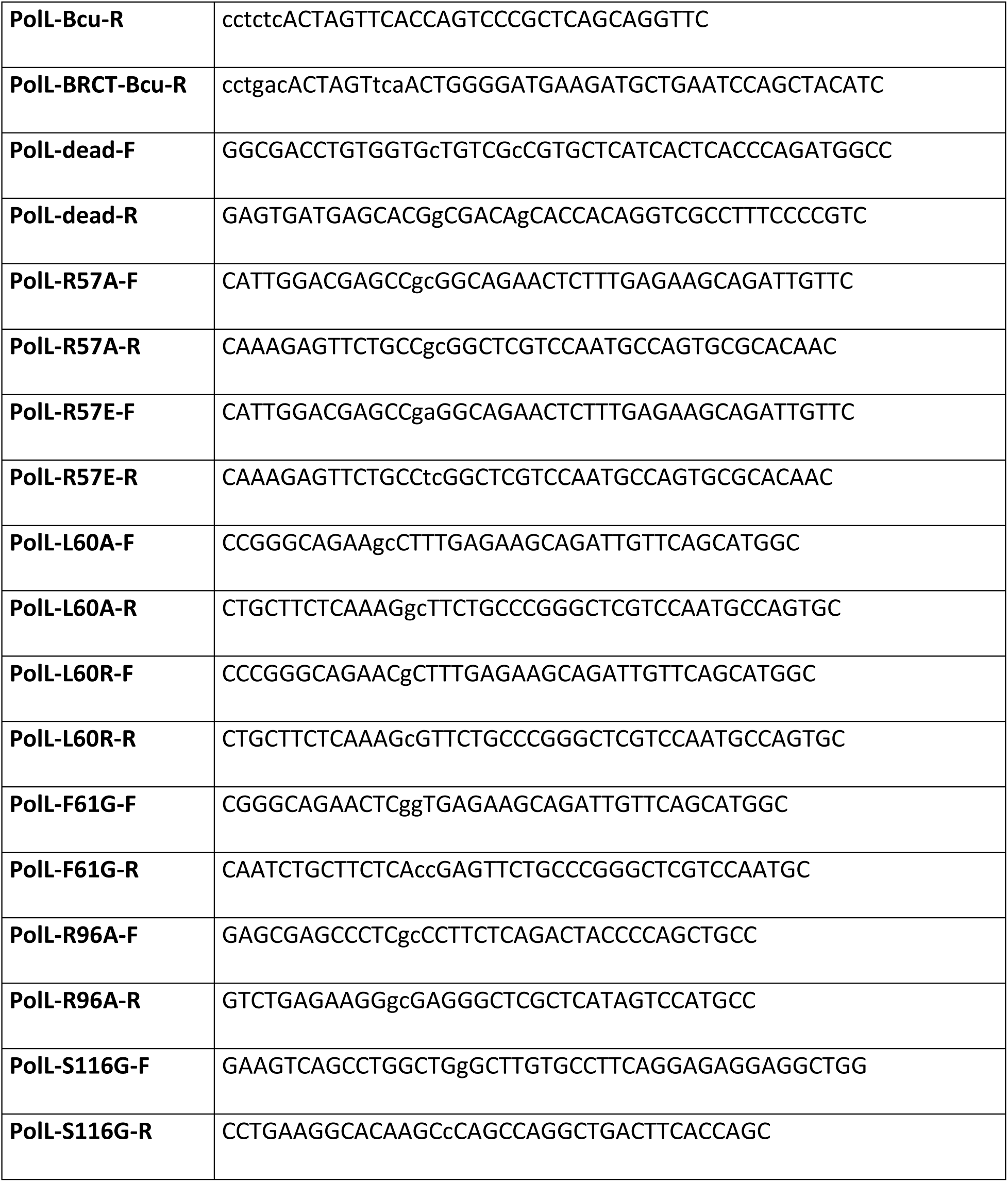

## Supplementary Information

**Table 1:**
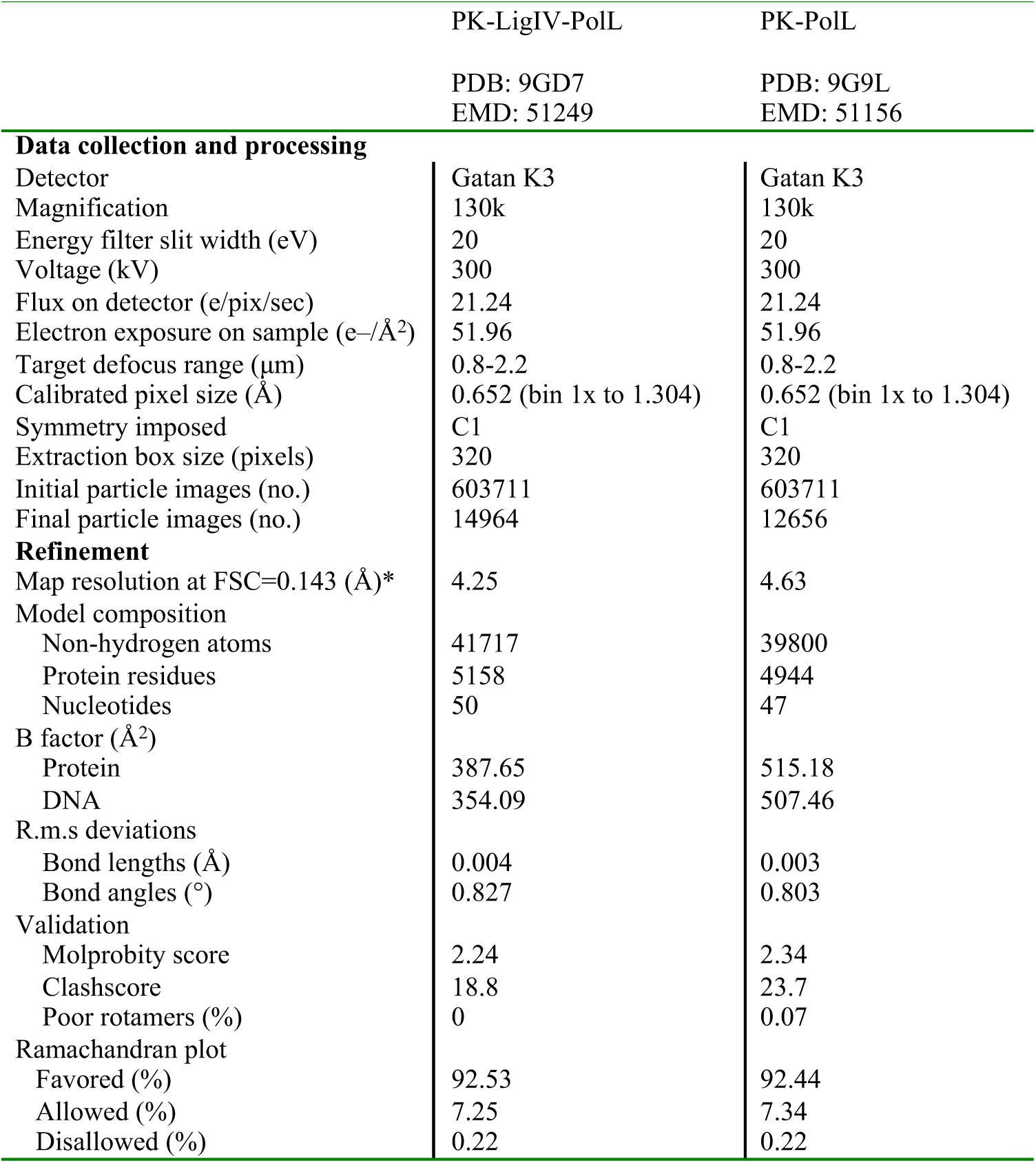
Cryo-EM data parameters and statistics.

**Figure S1:**
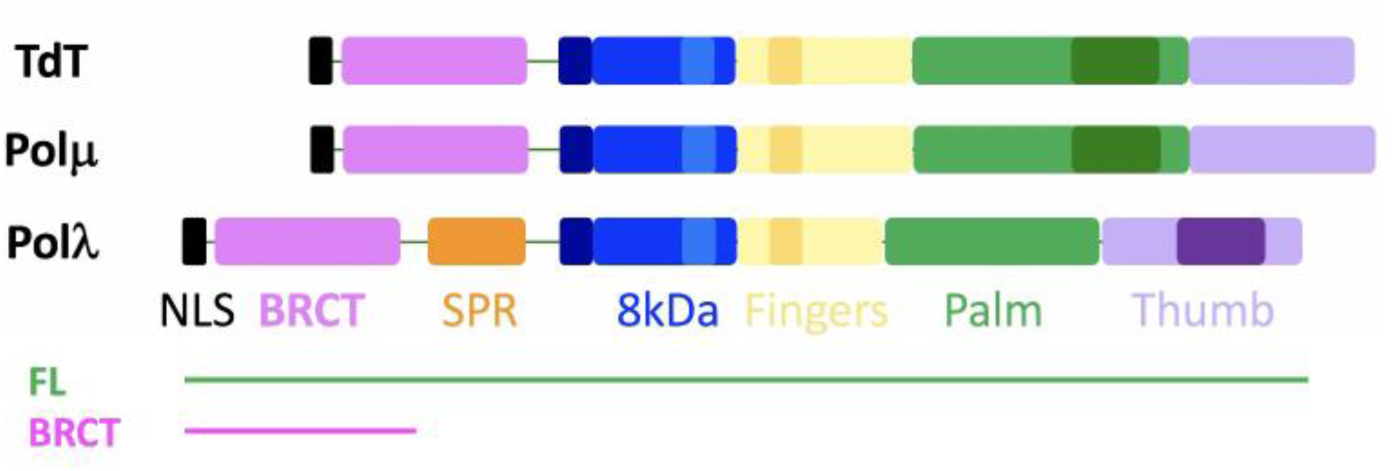
Domain organization of Pol X family DNA polymerases. The Pol λ constructs used in this study correspond to the full-length protein (FL) and the amino-terminal region (residues 1-136) containing the nuclear localization sequence (NLS) and the BRCT domain (BRCT).

**Figure S2:**
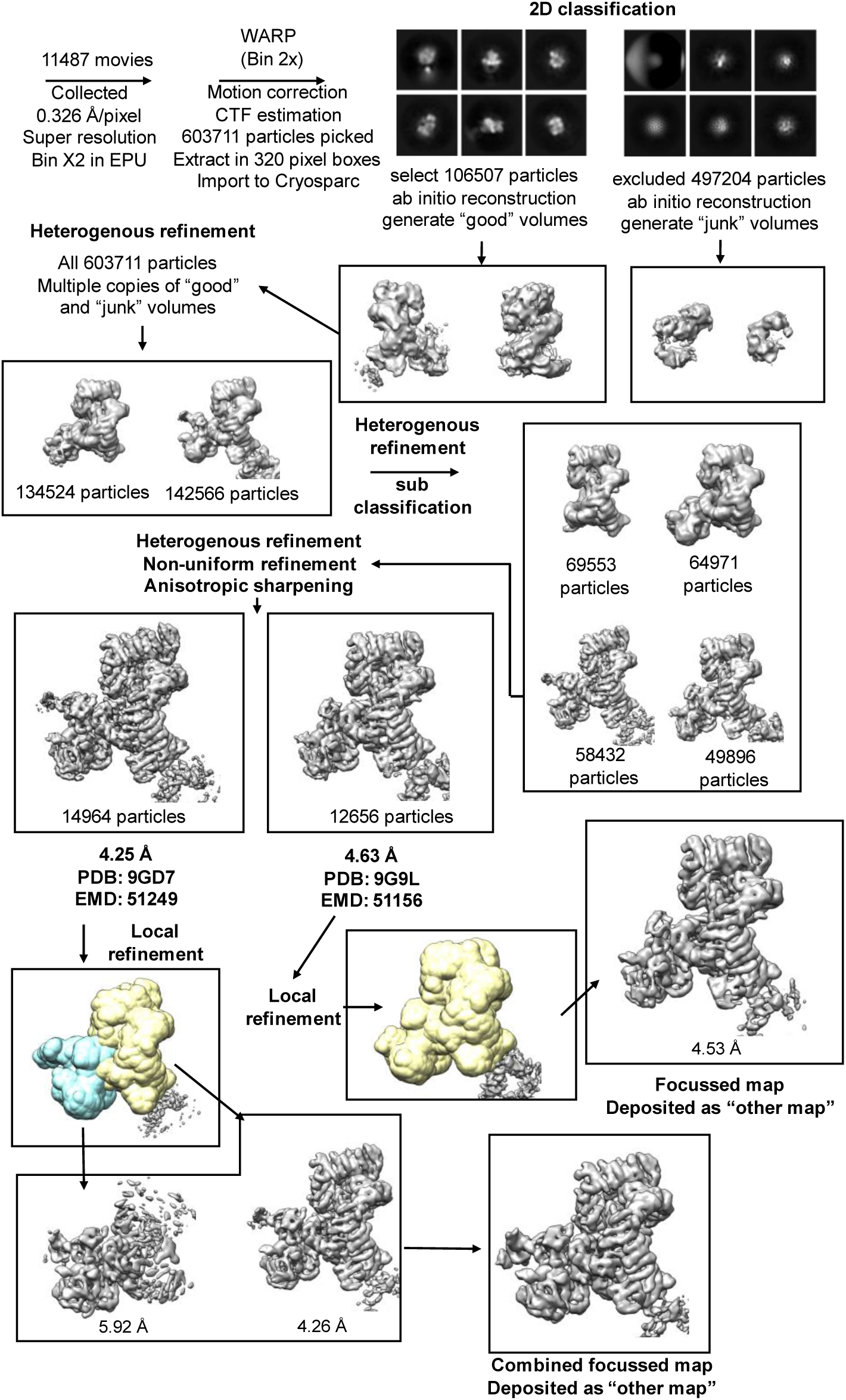
Single-particle cryo-EM image processing workflow for DNA-PK +PAXX + Pol λ with and without LX4. Schematic showing particle picking using WARP and processing including 2D classification and *ab initio* reconstruction using CryoSPARC. The two main classes generated with the corresponding number of particles is shown and the two maps following non-uniform refinement with resolutions for an FSC of 0.143 are given. Additional focused and composite maps are also shown.

**Figure S3:**
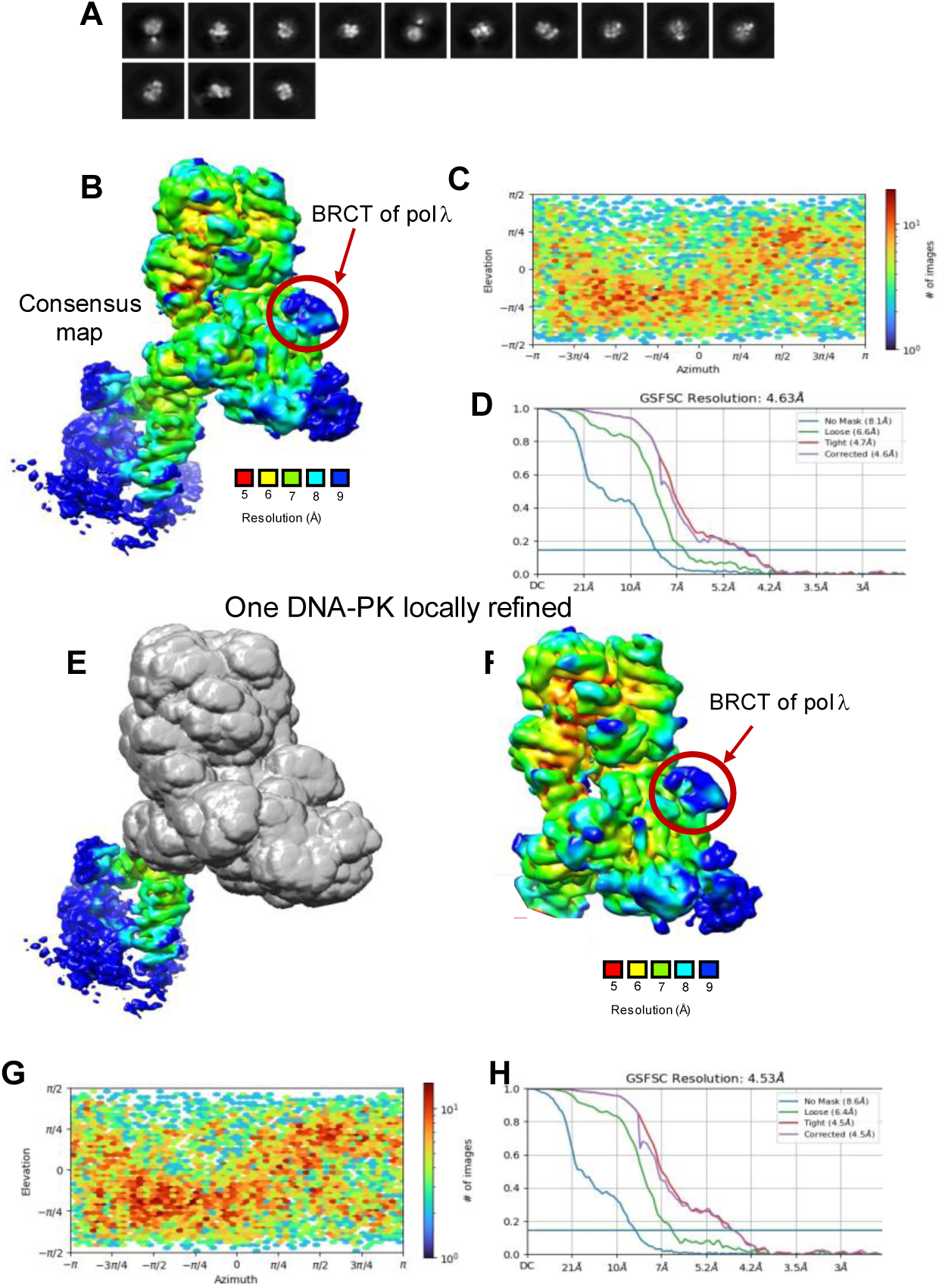
Cryo-EM data of DNA-PK with BRCT domain of Pol λ. **a)** Example of 2d classes. **b)** Local resolution map of DNA-PK dimer with BRCT domain of Pol λ consensus cryo-EM map. **c)** Angular distribution calculated in cryoSPARC for particle projections shown as a heat map of the consensus map. **d)** FSC resolution curves and viewing distribution plot of the consensus map. **e)** DNA-PK dimer with BRCT domain of Pol λ consensus cryo-EM map with masking area. **f)** Local resolution map of DNA-PK dimer with BRCT domain of Pol λ locally refined map. **g)** Angular distribution calculated in cryoSPARC for particle projections shown as a heat map of the locally refined map. **h)** FSC resolution curves and viewing distribution plot of the locally refined map. The colours corresponding to each resolution are displayed on the specific key chart below the maps.

**Figure S4:**
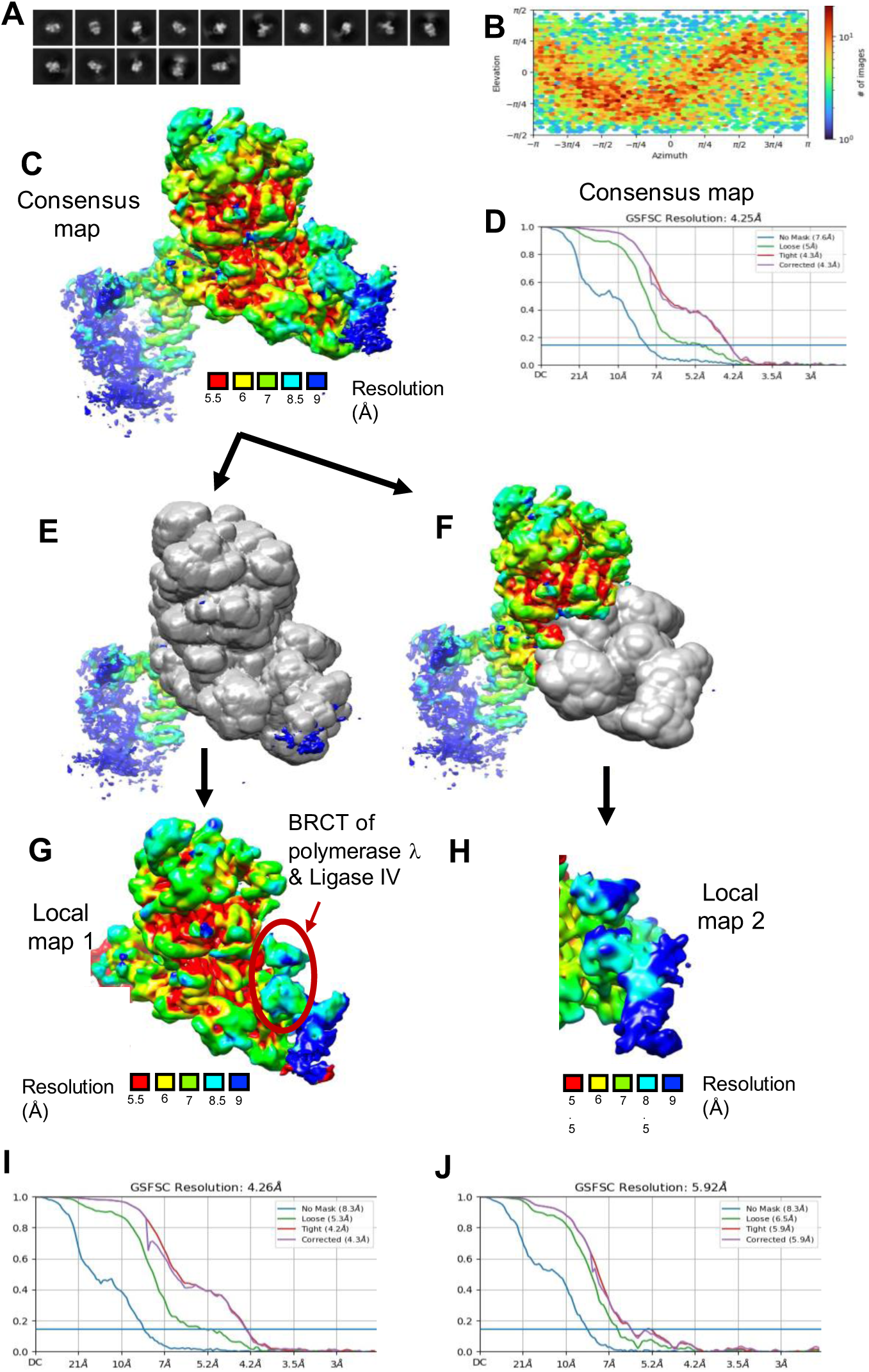
Cryo-EM data of DNA-PK with BRCT domain of Pol λ and LX4. **a)** Example of 2d classes. **b)** Angular distribution calculated in cryoSPARC for particle projections shown as a heat map of the consensus map. **c)** Local resolution map of DNA-PK dimer with BRCT domain of Pol λ and Ligase IV consensus cryo-EM map. **d)** FSC resolution curves and viewing distribution plot of the consensus map. **e)** and **f)** Consensus map with masking area 1 and 2, respectively. **g)** and **h)** Local resolution of the two locally refined maps. i) and **f)** FSC resolution curves and viewing distribution plot of local maps 1 and 2. The colours corresponding to each resolution are displayed on the specific key chart below the maps.

**Figure S5:**
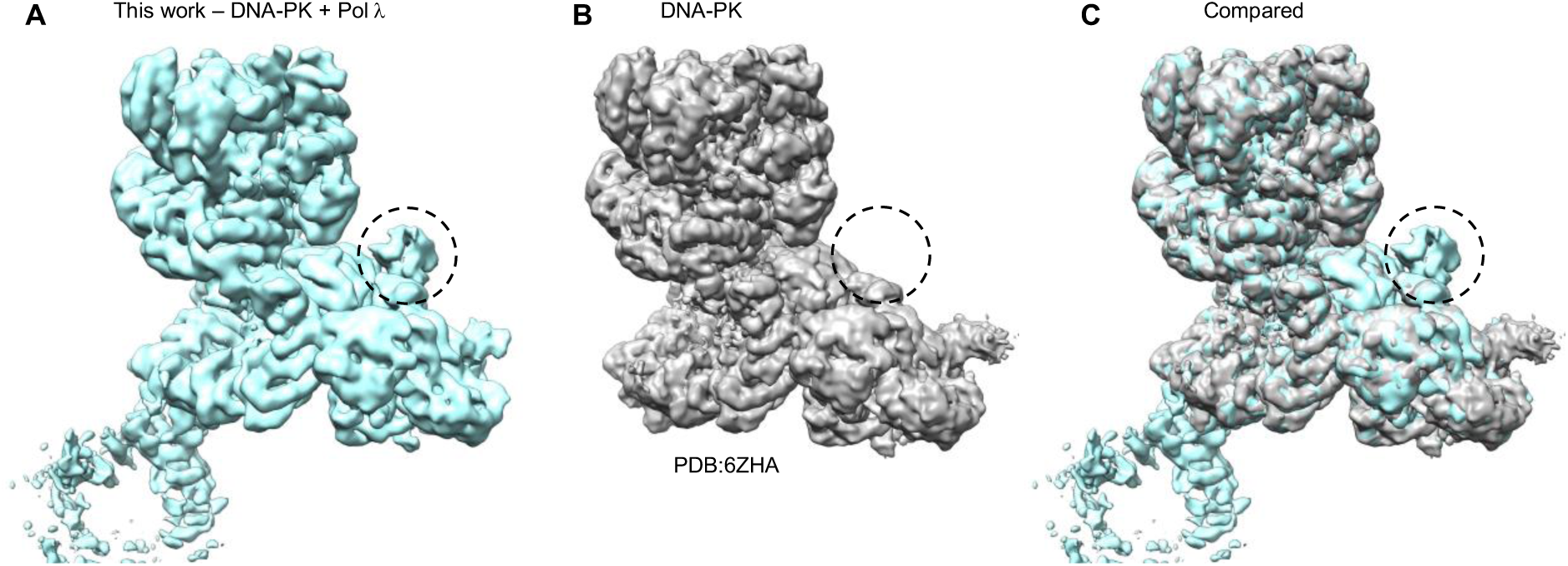
Comparison of cryo-EM maps. **A)** DNA-PK + Pol λ (this work) in blue. **B)** DNA-PK (PDB: 6ZHA), **C)** Comparison overlay of DNA-PK + Pol λ (blue) and DNA-PK (grey).

**Figure S6:**
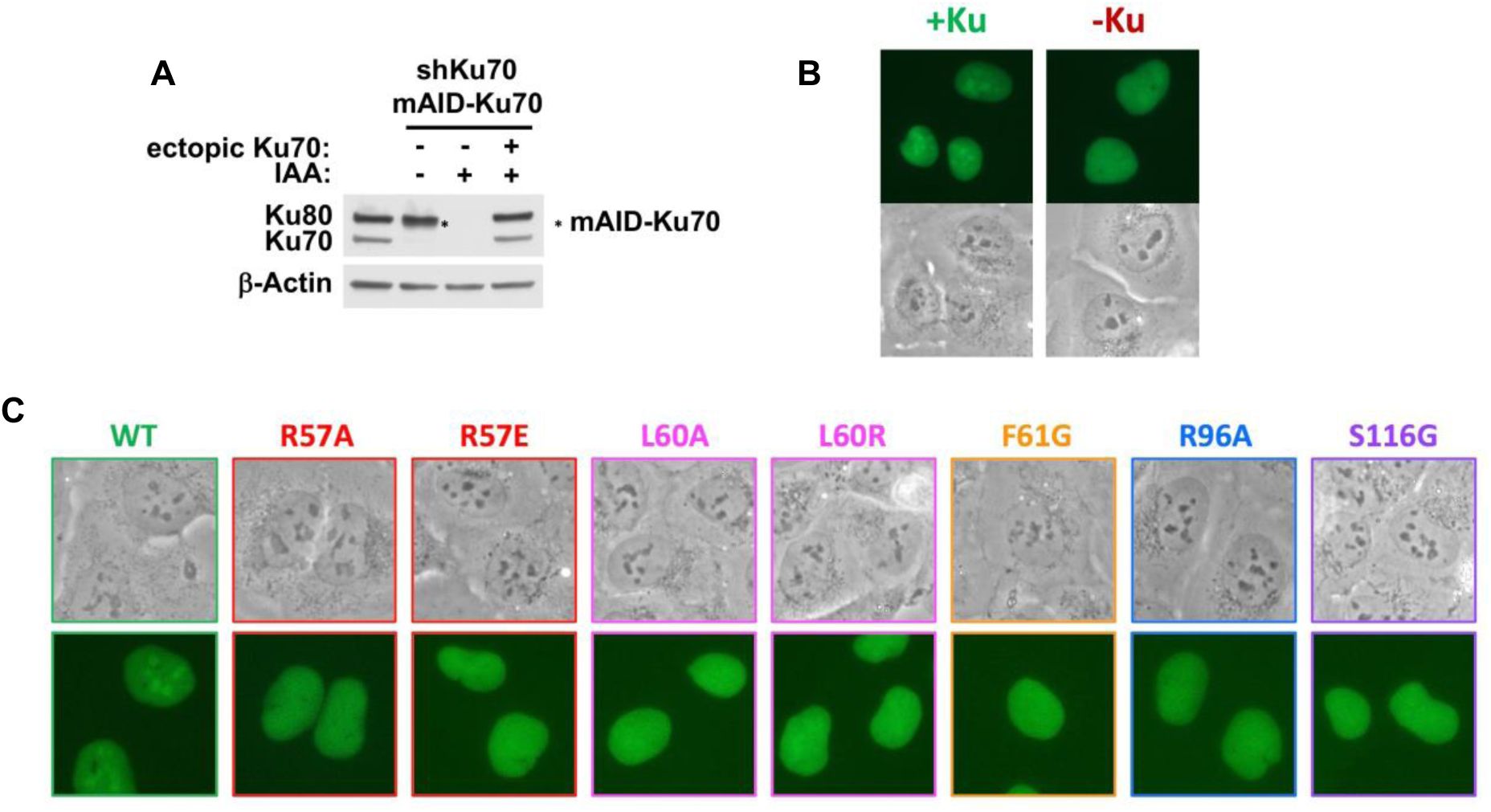
**(A)** Western blot on whole cell protein extracts from U2OS cells either unmodified (lane 1) or constitutively expressing an shRNA against endogenous Ku70 and rescued with expression of mAID-tagged Ku70 (lanes 2-4), treated or not with auxin (IAA) for 16 h. Asterisk indicates the position of mAID-Ku70 signal (MW: 77.4 kDa) below that of Ku80. **(B)** Fluorescence micrographs of U2OS/mAID-Ku70 cells (see (A)) expressing full-length GFP-tagged Pol l, in the presence (+Ku) or the absence (-Ku). **(C)** Fluorescence micrographs of U2OS cells expressing WT or mutated GFP-tagged Pol l BRCT domain.

**Figure S7:**
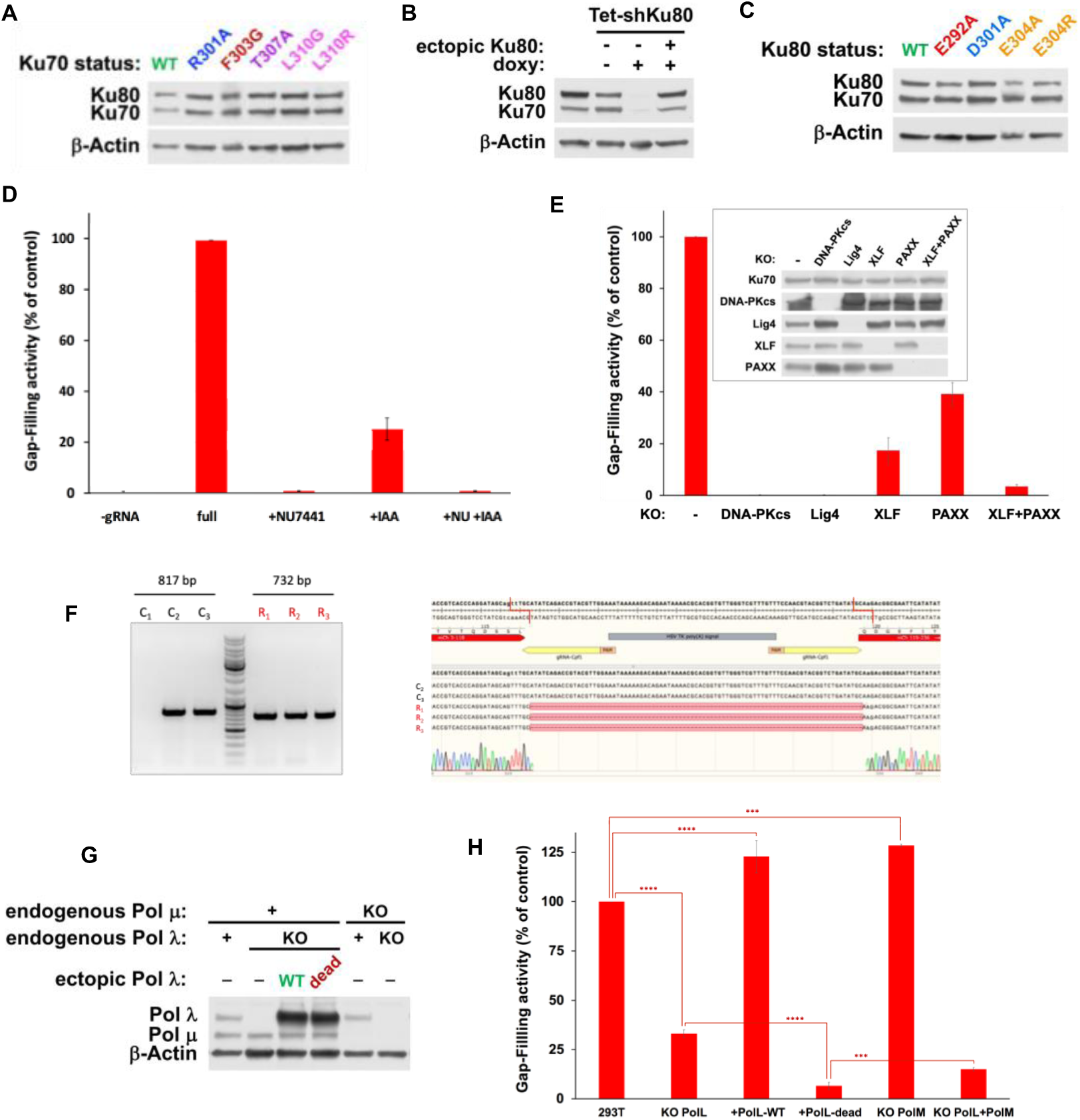
**(A)** Western blot on whole cell protein extracts from U2OS/mAID-Ku70 cells (see Suppl. Figure S6A) depleted of endogenous Ku70 and complemented with ectopic expression of either WT or mutated forms of Ku70, as indicated. **(B)** Western blot on whole cell protein extracts from U2OS cells either unmodified (lane 1) or expressing a doxycycline (doxy)-induced shRNA against endogenous Ku80 (lanes 2-4) and rescued with expression of Ku80 (lane4). **(C)** Western blot on whole cell protein extracts from U2OS/Tet-shKu80 cells (see (B)) depleted of endogenous Ku80 and complemented with ectopic expression of either WT or mutated forms of Ku80, as indicated **(D)** Gap-filling activity assessed in HEK-293T/mAID-Ku70 cells knocked-down for Ku70 when indicated (+IAA), in the presence or not of 3 µM DNA-PK inhibitor (+NU7441 or +NU). Results are normalized to the control condition (full) and plotted as mean values of five to thirteen experiments ± SD. **(E)** Gap-filling activity assessed in HEK-293T cells knocked-out (KO) for different NHEJ genes, as indicated. Results are normalized to the NHEJ-proficient parental HEK-293T cell line and plotted as mean values of four to five experiments ± SD. Inset: control western blot on whole cell protein extracts from the different KO cells. **(F)** DNA junction analysis following gap-filling assay. Left: agarose gel electrophoresis showing PCR products amplified around the junction following stable genome integration of the reporter substrate and gap-filling reaction (see the Materials and methods section). Three positive red-fluorescent clones (R1, R2 and R3) and three non-fluorescent clones as a control (C1, C2 and C3) were analyzed. Expected length of the PCR fragments are indicated. Right: the DNA sequences of the PCR fragments are aligned with the reporter substrate sequence using the Snapgene^®^ software (Dotmatics). **(G)** Western blot on whole cell protein extracts from HEK-293T cells knocked-out (KO) for POLL, POLM or both genes. When indicated, POLL KO cells were complemented with expression of ectopic WT or catalytic dead Pol λ. **(H)** Gap-filling activity assessed in HEK-293T cells knocked-out (KO) for POLL, POLM or both genes. Results are normalized to the parental HEK-293T cell line and plotted as mean values of four to fourteen experiments ± SD. P-values from Student’s t-test between the indicated conditions are as follows: 293T versus KO PolL (<0.0001 ****), 293T versus KO PolL + Poll-WT (<0.0001 ****), 293T versus KO PolM (0.0002 ***), KO PolL versus KO PolL + Poll-dead (<0.0001 ****), KO PolL+PolM versus KO PolL + Poll-dead (0.0001 ***).

**Figure S8:**
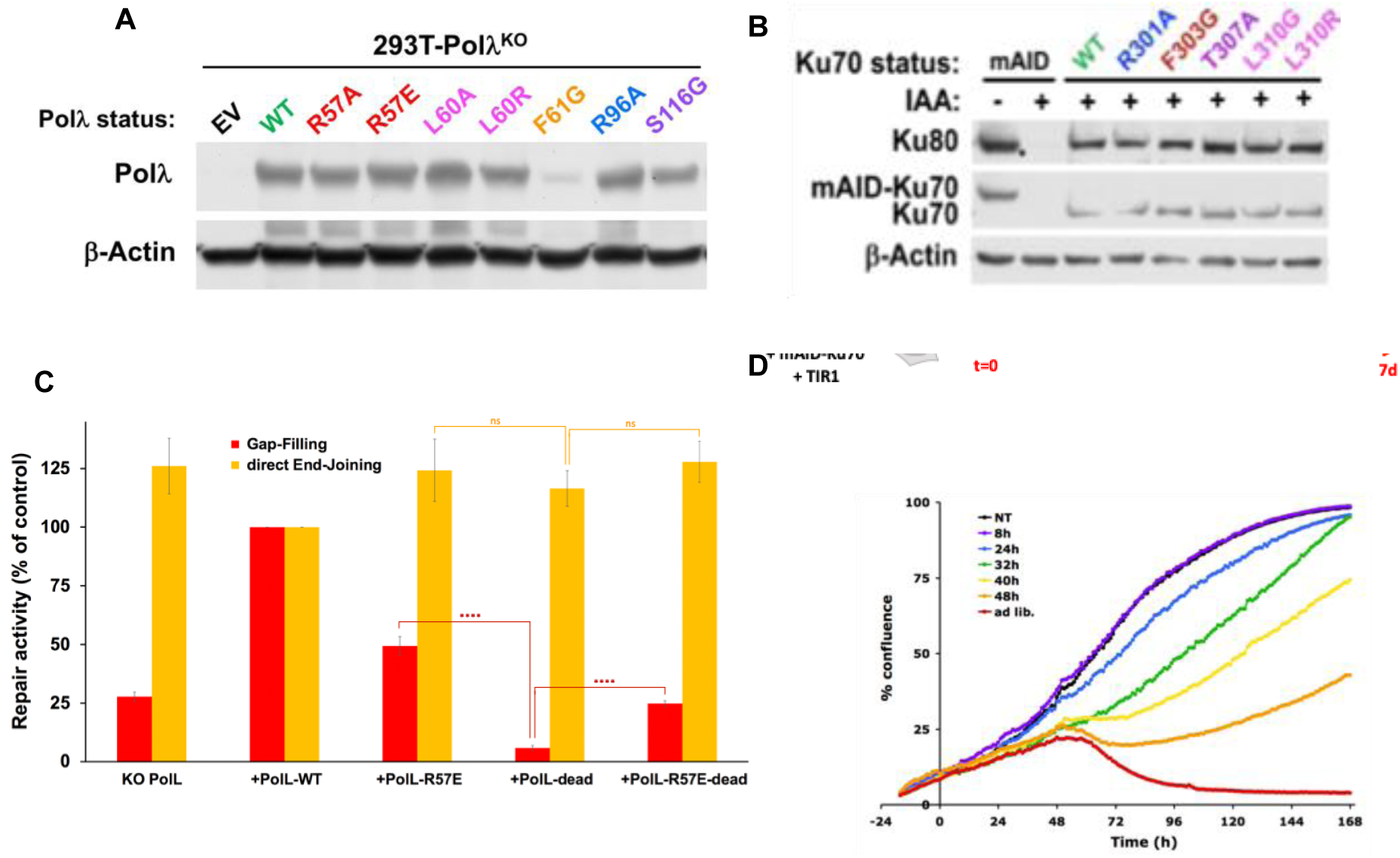
**(A)** estern blot on whole cell protein extracts from HEK-293T cells knocked-out (KO) for POLλ and complemented with an empty vector (EV) or expression vectors for wild-type (WT) or the indicated mutants of Pol λ. **(B)** Western blot on whole cell protein extracts from HEK-293T/mAID-Ku70 cells depleted of endogenous Ku70 in the presence of auxin (+IAA) and rescued with ectopic expression of either WT or mutated forms of Ku70, as indicated. Asterisk indicates the position of mAID-Ku70 signal just below that of Ku80, which persisted after previous hybridization of the membrane with anti-Ku70 antibody. **(C)** Gap-filling activity (red bars) or direct end-joining activity (orange bars) assessed in parallel in HEK-293T cells knocked-out for POLL and complemented with ectopic expression of wild-type (WT) or different mutants of Pol λ. Results are normalized to the WT condition and plotted as mean values of four to five experiments ± SD. P-values from Student’s t-test between the considered mutants, for gap-filling and direct end-joining activities, respectively, are as follows: Polλ-dead versus Poll-R57E (<0.0001 ****; 0.2862 ns), Poll-dead versus Poll-R57E-dead (<0.0001 ****; 0.0587 ns). **(D)** HEK-293T/mAID-Ku70 cells were seeded in 12-well plates and treated with auxin (+IAA) for the indicated time. Cell proliferation was then analyzed continuously up to 7 days by assessing confluence with an IncuCyte-ZOOM (Essen Bioscience).

